# KDM6A Loss Confers an Invasive Phenotype in Osteosarcoma by Activating Cytoplasmic YAP-Dependent β-catenin Stabilisation

**DOI:** 10.64898/2026.08.12.744422

**Authors:** Chinmay Nayak, Mukul Srivastava, Shibasish Chowdhury, Sudeshna Mukherjee, Rajdeep Chowdhury

## Abstract

Osteosarcoma (OS) is the most common primary malignant bone tumour and is characterised by aggressive growth, early metastasis, and a very stagnant clinical outcome. Although epigenetic dysregulation has been implicated in OS progression, the mechanisms linking epigenetic alterations to metastatic signalling remain unclear. Here, we identified the lysine-protein demethylase 6A (KDM6A/UTX) as a critical suppressor of OS metastasis and uncovered a novel regulatory axis involving the Hippo/YAP and Wnt/β-catenin signalling. Initial bioinformatics analyses revealed frequent KDM6A alterations and significantly reduced expression in OS patient datasets, which correlated with metastatic disease and poor prognosis. Functional inhibition of KDM6A by pharmacological inhibitors and siRNA induced a hyper-invasive phenotype, marked by elevated mesenchymal markers, enhanced cytoskeletal remodelling, increased transendothelial adhesion and decreased chemotherapeutic drug sensitivity. Importantly, restoration of KDM6A expression effectively counteracted these effects. Mechanistically, KDM6A loss activated Wnt/β-catenin signalling, resulting in nuclear translocation of β-catenin and transcriptional activation of genes associated with stemness and invasion. Therefore, inhibition of β-catenin reversed the invasive phenotype. Further analysis revealed that KDM6A regulated Hippo signalling through epigenetic control of the negative regulator of Yes-Associated Protein (YAP)-LATS1. KDM6A inhibition led to enrichment of H3K27me3, a repressive mark, at the LATS1 promoter. Accumulated YAP was predominantly localised in the cytoplasm, where it interacted with GSK3β and contributed to the stabilisation of β-catenin by preventing its proteasomal degradation. Collectively, our findings identify a novel KDM6A-LATS1-YAP-β-catenin signalling axis that drives metastatic progression in OS.

## 1. Introduction

Metastasis remains the leading cause of cancer-related mortality and represents one of the greatest challenges in modern oncology. The metastatic cascade requires cancer cells to acquire migratory, invasive, and stem cell-like properties that facilitate dissemination from the primary tumour and colonisation to distant organs (1). These phenotypic transitions are increasingly recognised as consequences of extensive transcriptional plasticity driven not only by genetic alterations but also by dynamic epigenetic reprogramming (2,3,4). In this regard, osteosarcoma (OS), the most common primary malignant bone tumour, is characterised by aggressive local growth, early metastatic dissemination, and poor clinical outcomes. The disease predominantly affects adolescents and young adults, with a second peak in incidence among older individuals. Despite substantial advances in surgical techniques and multimodal chemotherapy, the prognosis of patients with metastatic OS remains dismal, with five-year survival rates below 20%. Pulmonary metastasis remains the principal cause of mortality, underscoring the urgent need to identify molecular mechanisms that drive metastatic progression and uncover novel therapeutic vulnerabilities (5,6).

Among the diverse mechanisms that contribute to metastatic progression, epigenetic dysregulation has emerged as a critical determinant of tumour aggressiveness (2). Herein, histone methylation represents a highly dynamic and reversible epigenetic modification regulated by the opposing activities of histone methyltransferases (HMTs) and demethylases (KDMs), which together govern chromatin accessibility and transcriptional programs. In this context, the KDM family comprises several subclasses, including KDM1, KDM2, KDM3, KDM4, KDM5, KDM6, and KDM7, each targeting specific histone lysine residues and thereby regulating a broad spectrum of biological processes. Accumulating evidence suggests that KDMs exert highly context-dependent functions during tumorigenesis and cancer progression (7, 8). Depending on the cellular and molecular landscape, individual KDMs may act either as oncogenic drivers or tumour suppressors. For instance, KDM1A (LSD1) and members of the KDM4 family (KDM4A–C) are frequently overexpressed in multiple malignancies, including breast, prostate, and haematological cancers, where they promote tumour growth, metastatic dissemination, stemness, and immune evasion (9, 10). In contrast, KDM5C has been reported to function as a tumor suppressor in clear-cell renal cell carcinoma, highlighting the complex and context-specific nature of histone demethylase biology (11). Similarly, in OS, several KDM family members, such as KDM5 and KDM4, have been implicated in regulating cellular proliferation, stemness, chemoresistance, and metastatic potential (12, 13). These observations underscore the pivotal role of histone methylation/demethylation in OS progression, emphasising that the biological consequences of KDM activity are highly dependent on tumour context.

Among these enzymes, lysine demethylase 6A (KDM6A, also known as UTX) has emerged as a key epigenetic regulator of cellular differentiation and tumour suppression. KDM6A catalyses the removal of the repressive H3K27me3 mark, thereby promoting transcriptional activation of genes involved in lineage commitment and cellular homeostasis (14). Genetic alterations or loss of KDM6A have been reported in multiple malignancies and are frequently associated with poor clinical outcomes, increased tumour aggressiveness, and metastatic behaviour (15, 16). In a recent report, KDM6B was found to act as a tumour suppressor, and its downregulation led to greater pulmonary invasion (17). On the contrary, KDM6A inhibition increases sensitivity to anticancer drugs such as cisplatin (18). Nevertheless, the molecular mechanisms through which KDM6A regulates metastatic progression in OS remain poorly understood.

Importantly, according to a recent report, several signalling pathways implicated in OS progression are tightly regulated by epigenetic mechanisms. Among these, the Wnt/β-catenin and Hippo signalling pathways occupy central roles in controlling cell proliferation, differentiation, stemness, migration, and tissue regeneration (19). Aberrant activation of Wnt/β-catenin signalling is frequently observed in OS and has been linked to enhanced tumour growth, metastatic dissemination, and poor prognosis (20). Similarly, dysregulation of the Hippo pathway and activation of its downstream effector Yes-associated protein (YAP) promote tumour progression, therapeutic resistance, and metastatic competence (21, 22). Although these pathways have traditionally been studied independently, accumulating evidence suggests that β-catenin and YAP form an interconnected signalling network that coordinates transcriptional programs associated with cellular plasticity and metastasis. YAP can directly interact with components of the β-catenin destruction complex, regulate β-catenin stability, and cooperate with β-catenin-dependent transcriptional complexes to drive oncogenic gene expression (23, 24). Conversely, Wnt signalling can influence YAP activity through both canonical and non-canonical mechanisms. Such reciprocal regulation positions the YAP–β-catenin signalling node as a central integrator of oncogenic signals. Given their collective roles in promoting stemness, epithelial–mesenchymal transition (EMT)-like plasticity, invasion, and metastasis, dysregulation of the YAP–β-catenin axis may represent a critical signalling dependency in OS progression. However, the upstream epigenetic regulators that coordinate this signalling hub remain largely unknown.

In the present study, we identify KDM6A as a critical epigenetic gatekeeper of the YAP–β-catenin metastatic signalling axis in OS. We demonstrate that loss of KDM6A promotes a pronounced invasive phenotype through the activation of β-catenin signalling and non-canonical modulation of the Hippo pathway. KDM6A deficiency induces cytoplasmic retention of YAP, which enhances β-catenin stability and signalling output, thereby amplifying pro-metastatic transcriptional programs. Collectively, our findings uncover a previously unrecognised KDM6A–YAP–β-catenin regulatory network that drives advanced mesenchymal plasticity and invasive ability in OS, highlighting a potentially actionable signalling vulnerability for treating this aggressive disease.

## 2. Materials and Methods

### 2.1. *In Silico* Analysis of Publicly Available Datasets

Publicly available transcriptomic and genomic datasets were analysed to investigate the clinical and molecular relevance of KDM6A in OS. The Therapeutically Applicable Research to Generate Effective Treatments Osteosarcoma (TARGET-OS) dataset was accessed through the Genomic Data Commons (GDC) portal to assess the proportion of patients presenting with metastatic disease at diagnosis and to evaluate the association between metastatic status and overall survival (https://portal.gdc.cancer.gov/projects/TARGET-OS). Kaplan–Meier survival analysis was performed to compare the overall survival of patients with localised and metastatic disease. Furthermore, the GSE14359 dataset was analysed to identify differentially expressed histone demethylase genes in metastatic OS samples (https://www.ncbi.nlm.nih.gov/geo/query/acc.cgi?acc=gse14359). The transcriptomic data were analysed using the GEO2R tool under NCBI. In addition, the GSE86053 dataset was extracted to evaluate KDM6A expression in drug-tolerant persister cells compared with treatment-naïve cells. Pan-cancer genomic alteration analysis of KDM6A was performed using the cBioPortal by using Pan-cancer analysis of whole genomes (ICGC/TCGA, Nature 2020) (https://www.cbioportal.org/study/summary?id=pancan_pcawg_2020). The frequency and nature of KDM6A alterations across cancer types, the association between KDM6A alterations and overall survival, and the KDM6A alteration landscape in the bone cancer cohort were evaluated. Correlation analyses were also performed between KDM6A and components of the Hippo signalling pathway, including LATS1 and MOB1A, and between YAP and β-catenin, using the same dataset.

### 2.2. Chemicals and Reagents

TRI reagent (#T9424), fetal bovine serum (#F7524), RIPA buffer (#R0278), protease inhibitor cocktail (#P83409), Triton X-100 (#T8787), Tween 20 (#P1379-1L), Bradford reagent (#B6916), acrylamide (#A9099), N′,N′-methylenebisacrylamide (#M7279), GSKJ4 (#SML0701), chloroquine diphosphate salt (#C6628), Deferiprone (3-hydroxy-1,2-dimethyl-4(1H)-pyridone; #379409), and Verteporfin (#SML0534-5MG) were procured from Sigma-Aldrich. Trypsin (#25300062), Antibiotic-Antimycotic (100×; #15240062), and Opti-MEM (#31985070) were obtained from Gibco. DPBS (#21300025), Lipofectamine 3000 (#L3000015), DNase I, RNase-free (#EN-0521), Annexin V-FITC Conjugate (#A13199), and the MAGnify Chromatin Immunoprecipitation System (#492024) were purchased from Thermo Fisher Scientific. The iScript™ cDNA Synthesis Kit (#1708891), Clarity Max™ Western ECL Substrate (#1705062), Clarity Western ECL Substrate (#1705061), Immun-Blot PVDF Membrane (#1620177), and iTaq™ Universal SYBR® Green Supermix (#1725122) were procured from Bio-Rad. Glycine (#MB0131), Tris (#MB-029), sodium dodecyl sulfate (SDS; #MB010), skimmed milk powder (#GRM-1254), DMEM high glucose (#AL151A), McCoy’s 5A medium (#AL057A), MEM (#AL047S), glycerol (#GRM-081), dimethyl sulfoxide (DMSO; #TC185), bovine serum albumin (BSA; #MB083) and NP 40 (#MB143) were obtained from HiMedia. Pyrvinium pamoate (#HY-A0293) and Protein G Magnetic Beads (HY-KO204) was obtained from MedChemExpress and Luciferase Assay System (Promega). All primary antibodies used in this study were obtained from Cell Signalling Technology (CST), and the corresponding HRP-conjugated secondary antibodies were also procured from CST. siRNAs targeting the indicated genes and a scrambled control siRNA were obtained from Eurogentec. KDM6A overexpression vectors were obtained from Addgene. All other chemicals and reagents used in this study were of analytical grade or molecular biology grade, as appropriate.

### 2.3. Cell Culture

The human osteosarcoma cell lines (HOS and MG-63) and the rat osteosarcoma cell line UMR106 were procured from NCCS, Pune, and the EA.hy926 endothelial cells were a gift from Prof. Syamantak Majumder, Department of Biological Sciences, BITS Pilani, Pilani Campus. HOS and MG-63 were maintained in Minimal Essential Medium, whereas UMR106 and EA.hy926 cells were maintained in Dulbecco’s Modified Eagle Medium. All culture media were supplemented with 10% fetal bovine serum, and cells were cultured at 37°C in a humidified 5% CO atmosphere. Cells were routinely subcultured when they reached approximately 70–80% confluency.

### 2.4. Annexin V/PI Apoptosis Assay

Approximately 2 × 10^5^ cells were seeded in a 6-cm cell culture plate and allowed to grow overnight. Following the drug exposure, the culture media containing floating cells was transferred to a 15 mL Falcon tube. The adherent cells were detached by trypsinisation and collected in the same tube. The combined cell suspension was centrifuged at 2000 rpm, and the resulting pellet was washed once with 1X PBS, followed by a second centrifugation at the same speed. The obtained pellet was then resuspended in 500 μL of 1X binding buffer, after which annexin and propidium iodide (PI) were added according to the manufacturer’s instructions. All the samples were then incubated for 10 mins in dark, except the autofluorescence control. Apoptotic cell populations were then quantified using a CytoFLEX flow cytometer (Beckmann Coulter) with FITC-A (Green) and PE-A (red) detection channels. A total of 10,000 events were acquired for each sample, and the data were processed using CytExpert software.

### 2.5. Cell Viability Assay

Cell viability was determined using the MTT assay. Briefly, 6×10³ cells per well were seeded into 96-well plates and allowed to adhere overnight under standard culture conditions. Cells were subsequently treated with the indicated drugs for the specified duration. At the end of the treatment period, MTT reagent (0.5 mg/mL) was added directly to each well after removing the previous drug-containing media and the plates were incubated for 3h to allow formazan crystal formation. The crystals were then dissolved in DMSO, and absorbance was recorded at 570 nm with a reference wavelength of 630 nm using a Multiskan Sky microplate reader. Cell viability was calculated using the following formula.

% Cell viability = (Mean absorbance of treated cells / Mean absorbance of control cells) × 100

### 2.6. Pharmacological Inhibitors Used

KDM6A activity was inhibited pharmacologically using GSKJ4 (selective KDM6A inhibitor) and Deferiprone (pan KDM inhibitor) at a final concentration of 10 μM and 50 μM, respectively. β-catenin was inhibited using Pyrvinium pamoate (PP) at a final concentration of 50 nM. For YAP inhibition, cells were treated with Verteporfin (VP) at a final concentration of 5 μM. Unless otherwise specified, cells were treated with GSKJ4 for 24 h. For combination treatments, cells were pre-treated with the respective inhibitor for 6 h before the addition of GSKJ4. For proteasome inhibition experiments, cells were treated with MG132 at a final concentration of 0.5 μM. When MG132 was used in combination with VP, both compounds were administered for 3 h before GSKJ4 treatment. Chloroquine (10 μM) was used as an autophagy inhibitor, and when combined with VP, both compounds were administered for 3 h before GSKJ4 treatment.

### 2.7. siRNA Transfection

2-4 × 10^5^ cells were seeded in six-well culture plates and allowed to reach approximately 70% confluency. Cells were then transfected with siRNAs targeting KDM6A, YAP, or β-catenin using Lipofectamine 3000 reagent according to the manufacturer’s instructions. The designated siRNAs were used at the specified working concentrations. Scrambled siRNA was used as a negative control.

### 2.8. KDM6A Overexpression

For KDM6A overexpression experiments, 4 × 10^5^ cells were transfected with the KDM6A expression construct using Lipofectamine-based transfection according to the manufacturer’s instructions. Following transfection, cells were incubated under standard culture conditions and subsequently processed for immunoblotting and functional assays.

### 2.9. RNA Isolation, cDNA Synthesis, and qPCR

Total cellular RNA was extracted using TRI reagent. cDNA was synthesised using 2 μg of total RNA and subsequently used for quantitative real-time PCR analysis. Gene expression was quantified using the QuantStudio 3.0 Real-Time PCR System. β-Actin or GAPDH was used as the housekeeping control, and relative gene expression was calculated using the Pfaffl method. The primer sequences used in this study are presented in **Table 1**.

**Table 1.**
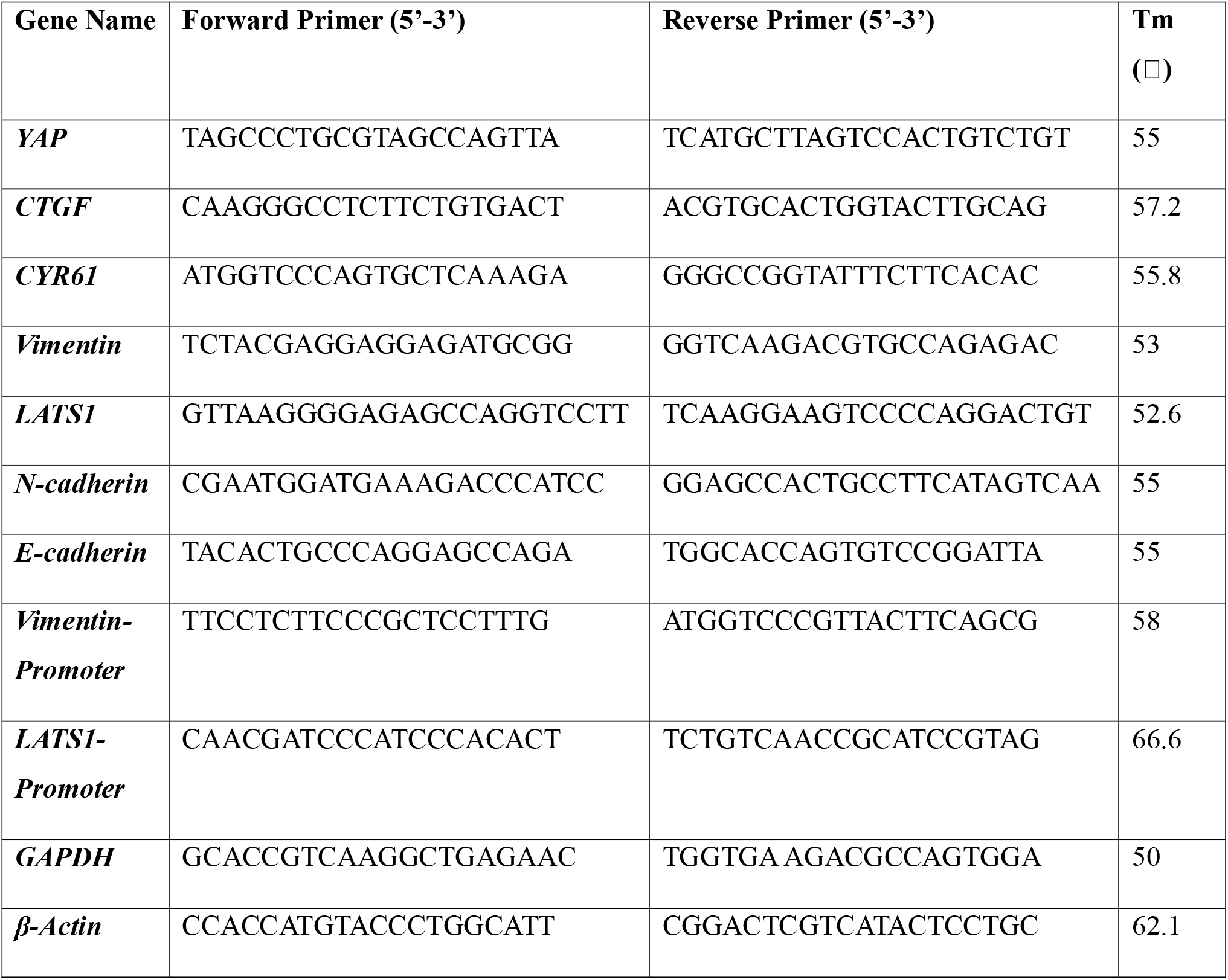
List of Primers Used in the Study.

### 2.10. Immunoblotting

Treated cells were lysed using a modified radioimmunoprecipitation assay (RIPA) buffer with added protease inhibitor. Total protein concentration was determined using the Bradford assay. Equal amounts of protein were mixed with 5X loading buffer and then denatured at 100 °C for 10 min. Protein samples were resolved by SDS-PAGE and subsequently transferred onto polyvinylidene difluoride (PVDF) membranes. Then the membranes were blocked with either 5% skim milk (for proteins except phospho-proteins) or 3% BSA (for phospho-proteins) in TBS, depending on the antibody used. The membranes were subsequently incubated with the respective primary antibodies, followed by incubation with appropriate HRP-conjugated secondary antibodies. Wherever required, membranes were stripped and re-incubated with additional primary antibodies. Protein bands were visualised using an enhanced chemiluminescence (ECL) detection system, and densitometric analysis was performed using ImageJ software. GAPDH, β-actin, or total H3 was used as the loading control, as required.

### 2.11. Immunofluorescence Analysis

For immunofluorescence staining, 2.5 × 10 cells were seeded onto sterile glass coverslips placed in 6-well culture plates. Once the desired cell confluency was achieved, cells were treated with the indicated drugs. Following the treatment, cells were washed with 1x PBS once and then fixed in 2% paraformaldehyde (PFA) for 10 min at room temperature. Fixed cells were washed 3x with 1x PBS, permeabilised with 0.1% Triton X-100 for 1-5 mins (depending on the target proteins), and then blocked with 2.5% bovine serum albumin (BSA) for 60 min at room temperature. Cells were then incubated overnight at 4**°C** with the targeted primary antibodies, which were diluted 1:1000 in 2.5% BSA. Subsequently, excess primary antibody was removed by washing with PBS, after which Alexa Fluor-conjugated secondary antibodies, which were diluted at 1:2000 in 2.5% BSA, were applied for 90 min at room temperature. DAPI was used to stain the nucleus. For F-actin staining, cells were labelled with green fluorescent phalloidin, followed by DAPI staining. The stained coverslips were mounted in 70% glycerol and imaged using the Zeiss Axio Observer Z1/7 Apotome microscope.

### 2.12. Wound-Healing Assay

Cells were seeded in six-well culture plates and allowed to reach 90 % confluence. A uniform linear scratch was generated across the cell monolayer using a 200 μL sterile pipette tip. The wells were gently washed with PBS to remove detached cells and replenished with fresh culture medium containing the indicated treatments. Images were acquired immediately after scratching (0 hr) and at the indicated time points using an inverted microscope. The percentage of wound closure was quantified using ImageJ software.

### 2.13. Transwell Migration Assay

The cell migration assay was performed using the Transwell chambers. Following the indicated treatments, cells were suspended in serum-free medium and seeded into the upper chamber of the transwell inserts. The lower chamber was filled with complete culture medium containing serum as a chemoattractant. Cells were incubated under standard culture conditions for 12 hours to allow migration through the membrane. Non-migrated cells were removed from the upper surface of the membrane, whereas migrated cells on the lower surface were fixed and stained with crystal violet. Images from multiple fields were acquired using an inverted microscope with Zen 2.3 SP1 software, and the number of migrated cells was quantified across multiple microscopic fields using ImageJ.

### 2.14. Transendothelial Adhesion Assay

For the hetero-adhesion assay, EA.hy926 endothelial cells were cultured until a confluent monolayer formed. Following the indicated treatment, OS cells were stained with DiI dye, added to the endothelial monolayer, and incubated for 30 mins to allow cell–endothelial adhesion. The wells were subsequently washed gently with PBS to remove non-adherent cells and fixed with 4% PFA. Adherent OS cells were visualised using a fluorescence microscope (Zeiss Axio Observer Z1/7 Apotome), and Images from multiple fields were acquired. Image acquisition and analysis were performed using Zen 2.3 SP1 and ImageJ, respectively.

### 2.15. Static Cell-Matrix Adhesion Assay

For the static adhesion assay, culture plates were first coated with 1 % gelatin and incubated in a CO_2_ incubator for 2 hours before adhesion. Following the indicated treatment time points, OS cells were seeded onto the gelatin-coated surface and allowed to adhere. Non-adherent cells were removed by gentle washing with PBS. The remaining adherent cells were stained with crystal violet. Then the plates were visualised by an inverted microscope and quantified using ImageJ.

### 2.16. Spheroid Formation and Invasion Assay

The spheroid formation and invasion assay was performed following the earlier reported method with some modifications (21). HOS cells were suspended in MEM supplemented with 0.2% methylcellulose at an appropriate cell density. Hanging drops (10 μL) of the cell suspension were generated and incubated at 37 °C under 5% CO_2_ for 3 days to allow spheroid formation. To prevent evaporation and drying of the hanging drops, tissue culture plates containing sufficient PBS were used during incubation. After spheroid formation, individual spheroids were carefully transferred to 96-well round-bottom plates containing 100 μL of the complete culture medium and incubated for 24 h to allow acclimatisation. Following acclimatisation, spheroids were treated with the indicated drugs by adding culture medium to a final volume of 200 μL per well and incubated for 4 days. Phase-contrast images of spheroids were captured every 2 days, starting from days 0 through 6. Spheroid area and perimeter were quantified and compared with those of untreated control spheroids. Image analysis was performed using ImageJ, and line plots were generated using GraphPad Prism.

For the 3D spheroid invasion assay, 96-well round-bottom plates were first coated with 1% gelatin and incubated in a CO2 incubator for 2 hours before spheroids were transferred onto them. The extent of spheroid invasion was assessed by measuring the radial outgrowth from the spheroid body. Phase-contrast images of spheroids were captured every 2 days, starting from days 0 through 4

### 2.17. Cellular Fractionation

Following the treatments, cells were harvested and subjected to subcellular fractionation to obtain cytoplasmic and nuclear protein fractions. Cells were harvested and washed thoroughly with PBS. The cell pellet was resuspended in ice-cold mild lysis buffer [NP-40, Protease Inhibitor Cocktail. The samples were then centrifuged for 10 minutes at 10,000 rpm, and the nuclear fraction was obtained as a pellet, whereas the supernatant was collected as the cytoplasmic fraction. Furthermore, the nuclear lysate was obtained by resuspending the nuclear pellet in harsh lysis buffer. Cells were further lysed with sonication (15 kW, 2 cycles, 15sec each), followed by immunoblotting. GAPDH and total H3 were used as cytoplasmic and nuclear fraction controls, respectively.

### 2.18. Chromatin Immunoprecipitation–Quantitative PCR

Chromatin immunoprecipitation (ChIP) was performed using the MAGnify immunoprecipitation system. For this, cells were cultured in 10-cm culture plates and treated with drugs for the specified duration. Following treatment, cells were harvested and washed with PBS. Protein–DNA complexes were cross-linked using 1% formaldehyde and subsequently neutralised using 1.25 M glycine. Following additional PBS washes, cells were lysed in lysis buffer containing a protease inhibitor cocktail (50 μL per 10^6^ cells). Chromatin was fragmented by sonication for 60 cycles (45s on, 15 s off), and the efficiency of chromatin shearing was confirmed by agarose gel electrophoresis. Fragmented chromatin was incubated overnight with the respective antibodies against H3K27me3 and β-catenin at a dilution of 1:50. Following immunoprecipitation, immune complexes were washed, eluted, and reverse cross-linked. DNA was purified following proteinase K digestion. Enrichment of the LATS1 and Vimentin promoter regions was quantified by qPCR using gene-specific primers. The ChIP assay was performed following the earlier reported protocol (21).

### 2.19. Luciferase Assay

The Luciferase assay was performed following the earlier reported method with some modifications (21). Cells were seeded at a density of 2.5 × 10^5^ cells per well in six-well plates and transfected with the 8×GTIIC-luciferase reporter plasmid using the Lipofectamine reagent. After 6 h of transfection, the medium was replaced, and cells were treated with GSKJ4 for the indicated duration. Luciferase activity was assessed using Promega’s Luciferase Assay Kit according to the manufacturer’s protocol. Briefly, cells were washed with 1x PBS and lysed using the lysis buffer provided in the kit. Lysates were collected by scraping and centrifuged at 2500 rpm for 15 min. Equal volumes of lysate were mixed with 100 μL of Luciferase Assay Reagent, and luminescence was immediately measured using the GloMax 20/20 luminometer (Promega).

### 2.20. Co-Immunoprecipitation Assay

Whole cell protein lysate was prepared as described above. For each immunoprecipitation reaction, 500 μg of total protein was incubated with Protein G Magnetic Beads, which were pre-bounded to either anti-YAP, anti-β-catenin and negative control anti-rabbit IgG. Antibody binding was carried out by rotating the bead suspension at 4 °C for 4 h, followed by incubation with the protein lysate overnight at 4 °C on a rota spin. The following day, immune complexes were recovered using the magnetic beads, washed repeatedly with PBS to remove non-specifically bound proteins and subjected to SDS-PAGE together with the corresponding input samples. Protein-protein interaction involving YAP, β-catenin, and their binding partners was subsequently detected by Western blot using the appropriate primary and secondary antibodies.

### 2.21. In Vivo Lung Metastasis Model

Male Sprague Dawley (SD) rats aged 4-6 weeks with a body weight of 60±10 g were procured from the central animal facility at BITS Pilani. All animal procedures were approved by the Institutional Animal Ethics Committee (IAEC) at the Department of Pharmacy, BITS Pilani, Pilani campus (Protocol Number: IAEC/RES/38/04). Animals were housed under controlled laboratory conditions with a 12h light/dark cycle with access to adequate food and water.

Before the start of the experiment, animals were randomly divided into two experimental groups (n=5 per group). UMR106 OS cells were cultured under standard conditions and treated with GSKJ4 for 24 hours before injection. After the treatment, cells were harvested during the logarithmic growth phase, washed twice with sterile PBS, and resuspended in sterile PBS to form the cell suspension. Cell viability was confirmed to be greater than 90% before injection using the trypan blue method. Each rat received around 10^7^ UMR106 cells *via* lateral tail vein injection. The rats in the control group were injected with untreated UMR106 cells, whereas the second group received GSKJ4-pretreated UMR106 cells. Post-tumour cell injection, rats in the second experimental group were administered GSKJ4 at a dose of 4mg/kg body weight by intraperitoneal injection on alternate days for 20 days, while the control group received an equivalent amount of PBS according to the same treatment schedule. Throughout the experimental period, the animals’ body weight and any indications of distress were tracked. On day 22 after tumour cell injection, all animals were euthanised, and the lungs were carefully excised for assessment of metastatic burden. Gross metastatic nodules visible on the lung surface were counted, after which the lung tissues were fixed in 10% neutral-buffered formalin, paraffin-embedded, sectioned, and stained with hematoxylin and eosin (H&E) for histopathological examination. Pulmonary metastatic burden was evaluated by quantifying the number of gross metastatic nodules and histologically confirmed metastatic lesions in H&E-stained lung sections.

### 2.21. Statistical Analysis

Statistical analyses were performed using GraphPad Prism software, version 8.0. Depending on the experimental design, either an unpaired two-tailed Student’s t-test or one-way analysis of variance (ANOVA) was used to assess the significance of differences between treatment groups. For multiple comparisons, Tukey’s test was employed. Unless otherwise indicated, all data are presented as mean ± SD from at least three independent biological replicates.

Statistical significance was defined as p ≤ 0.05 and represented as follows: *p ≤ 0.05, **p ≤ 0.01, and ***p ≤ 0.001.

## 3. Results

### 3.1. KDM6A functions as a suppressor of advanced mesenchymal phenotype in OS

OS is the most common primary malignant bone tumour and is characterised by aggressive local growth and a strong propensity for distant metastasis. To assess the clinical burden of metastatic disease, we analysed OS patient data from the Therapeutically Applicable Research to Generate Effective Treatments Osteosarcoma (TARGET-OS) cohort available through the Genomic Data Commons (GDC) portal. Our analysis revealed that approximately 30% of patients present metastatic disease at the time of diagnosis **(Fig. 1A)**. Consistent with above, the presence of metastasis was associated with a markedly poorer prognosis, as evidenced by the Kaplan–Meier survival plot demonstrating a substantial reduction in 5-year overall survival compared with patients with localized disease **(Fig. 1B)**. Metastatic dissemination is a highly dynamic and multistep process that requires tumor cells to acquire phenotypic plasticity. This process is frequently accompanied by reversible alterations in epithelial and mesenchymal gene expression programs, reflecting transitions between distinct cellular states (25). Such plasticity is increasingly recognised as governed by epigenetic mechanisms that facilitate context-dependent transcriptional reprogramming rather than by stable mutations (26). These observations suggest that epigenetic regulators may play critical roles in orchestrating metastatic progression in OS. To gain mechanistic insight into the molecular drivers of OS metastasis and to investigate their potential connection to epigenetic regulation, we systematically examined the expression landscape of histone demethylase genes using transcriptomic data from the publicly available GSE14359 cohort. Given the established role of histone demethylases in modulating chromatin accessibility and transcriptional programs, we hypothesised that dysregulation of specific members of this family may contribute to metastatic competence. Notably, this analysis identified KDM6A as one of the most significantly downregulated epigenetic regulators in lung metastasised OS, implicating its potential involvement in disease progression and metastatic dissemination as well **(Fig. 1C)**. Given the established role of KDM6A in regulating chromatin architecture and transcriptional programs through demethylation of repressive histone marks, we hypothesized that its reduced expression may contribute to the acquisition of a more permissive epigenetic state that facilitates tumor cell plasticity and metastatic competence. Therefore, to further explore the relevance of KDM6A in therapy-associated cellular adaptation, we performed an independent analysis (GEO Series Accession # GSE86053) of transcriptomic datasets from drug-tolerant persister (DTP) cells (28, 43). These persister cells represent a transient, reversible cellular state characterised by extensive epigenetic reprogramming and enhanced survival under therapeutic stress. Intriguingly, KDM6A expression was markedly reduced in the persister cell population (OS-P) compared with treatment-naïve counterparts **(Supplementary Fig. 1A)**. Hence, the observed downregulation of KDM6A in both metastatic OS samples and DTP points to a potential relevance of KDM6A in metastatic progression and therapy tolerance, raising the possibility that loss of KDM6A-mediated chromatin regulation may promote adaptive transcriptional states supporting tumour persistence, dissemination, and treatment resistance.

**Figure 1.**
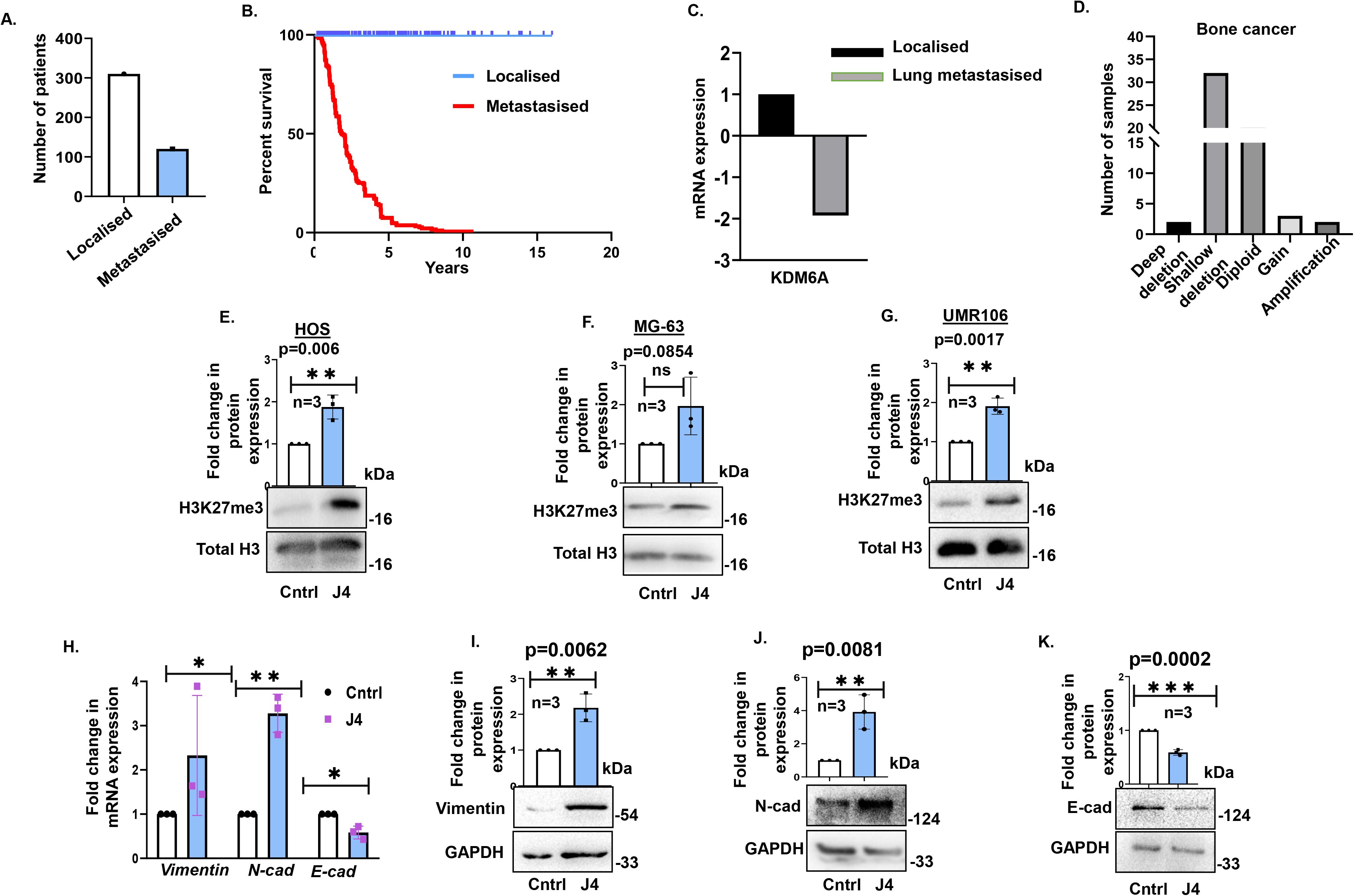
KDM6A is downregulated in OS, and its inhibition promotes an invasive phenotype. **(A)** Number of cases with localised OS or with metastasis at the time of diagnosis as obtained from TARGET-OS. **(B)** Kaplan-Meier survival analysis from TARGET-OS showing the percentage survival of OS patients with or without metastasis at diagnosis. **(C)** mRNA expression of KDM6A in metastasised compared to localised OS as obtained from the GSE14359 cohort GEO database. **(D)** Relative abundance of KDM6A genomic aberrations in bone cancer samples as obtained from cBioPortal. **(E)** Immunoblot analysis showing H3K27me3 expression in untreated HOS cells (control; Cntrl) or upon GSKJ4 (J4) treatment; total H3 served as the loading control. **(F)** Immunoblot analysis showing H3K27me3 expression in MG63 cells under control versus J4 conditions; total H3 served as the loading control. **(G)** Immunoblot analysis showing H3K27me3 expression in UMR106 cells upon J4 exposure; total H3 served as the loading control. **(H)** Fold change in mRNA expression of Vimentin, N-cad and E-cad in J4-treated cells. Immunoblot analysis showing Vimentin **(I),** N-cadherin **(J)** and E-cadherin **(K)** expression in J4-treated cells; GAPDH is the loading control. Unless otherwise specified, treatments were conducted for 24 h, and cells were treated with the KDM6A inhibitor J4 (10 μM). Fold-change values are expressed relative to the control group, which was set to 1. All data are presented as mean ± SD from at least three independent biological replicates (n = 3). Statistical significance between two groups was determined using an unpaired two-tailed Student’s t-test. Statistical significance is indicated as (*) p<0.05, (**) p<0.005 and (***) p<0.005. Cntrl denotes untreated cells. J4, GSKJ4; KDM6A, Lysine Demethylase 6A; N-cad, Neural cadherin; E-cad, Epithelial cadherin; GAPDH, Glyceraldehyde 3-phosphate dehydrogenase.

To characterise the genomic alteration landscape of KDM6A across human cancers, we performed a pan-cancer *in silico* analysis using the cBioPortal database. Analysis of the cBioPortal pan-cancer dataset revealed that KDM6A is frequently altered across multiple cancer types, with frequent shallow deletion (hemizygous state) representing the predominant event **(Supplementary Fig. 1B)**. Notably, Kaplan–Meier survival analysis her demonstrated that pan cancer patients harbouring KDM6A alterations exhibited significantly poorer overall survival compared with those lacking such alterations, highlighting the potential clinical relevance of KDM6A dysregulation **(Supplementary Fig. 1C)**. Likewise, analysis of the bone cancer cohort within the cBioPortal pan-cancer dataset also reflected similar trend of KDM6A shallow deletion than copy number gain, supporting a potential role for KDM6A loss in OS pathogenesis **(Fig. 1D)**. Given the *in-silico* findings, we sought to experimentally determine whether loss of KDM6A contributes to more invasive features. To address this, KDM6A was inhibited in three OS cell lines (HOS, MG-63, and UMR 106) using selective or pan pharmacological inhibitors (GSKJ4 and Deferiprone) or siRNA-mediated knockdown. Successful inhibition of KDM6A was confirmed by assessing H3K27me3 levels, a substrate of the KDM6 family. As expected, KDM6A inhibition resulted in a marked increase in H3K27me3 levels **(Fig. 1E–G; Supplementary Fig. 1D–E).** Interestingly, suppression of KDM6A consistently promoted expression of mesenchymal markers such as N-cad and Vimentin, along with a concomitant reduction in the epithelial marker E-cad at both the transcript (**Fig. 1H)** and protein levels in HOS cells **(Fig. 1I-1K & Supplementary Fig. F-G)**. To further validate these findings, KDM6A was inhibited in two additional OS cell lines, MG-63 and UMR106. Consistent with our earlier observations, KDM6A inhibition in both cell lines increased mesenchymal and decreased epithelial marker expression **(Supplementary Fig. 2A-2D)**. As an alternative approach, overexpression of KDM6A led to significant downregulation of N-cadherin and Vimentin in HOS cells **(Supplementary Fig. 2E-2I**). To further validate these findings, KDM6A was also overexpressed in MG-63 cells, which resulted in the upregulation of E-cadherin and downregulation of Vimentin **(Supplementary Fig. 2J-2M)**. During invasion, cancer cells undergo a phenotypic transition marked by the acquisition of mesenchymal features. This transition has been linked to increased aggressiveness, drug resistance, and cancer stemness (44). Our results identify KDM6A loss as a key driver of a more aggressive mesenchymal phenotype in OS.

### 3.2. Acquisition of enhanced mesenchymal features confers increased invasive capacity in OS cells

Upon confirming that KDM6A inhibition potentiates increased mesenchymal features in OS cells, we monitored the overall cytoskeletal remodelling of these cells following KDM6A inhibition. This resulted in a significant alteration of F-actin dynamics, as evidenced by increased phalloidin staining **(Fig. 2A)**, suggesting a role for KDM6A in the regulation of actin cytoskeletal organisation. To further evaluate the migratory capacity of the cells, a scratch assay was performed. Pharmacological inhibition of KDM6A with J4 significantly enhanced cell proliferation and migration **(Fig. 2B)**, whereas KDM6A overexpression markedly suppressed wound-healing capacity **(Supplementary Fig. 3A)**. Then to substantiate these observations, we also performed a transwell migration assay to assess invasiveness. Consistent with the scratch assay results, inhibition of KDM6A significantly increased the migratory capacity of the HOS cells, as evident by the increased number of cells migrating through the transwell upon KDM6A inhibition (**Fig. 2C).** Moreover, to understand whether the enhanced migratory phenotype observed following KDM6A inhibition correlates with increased invasive and metastatic potential, we executed a series of functional assays, including trans-endothelial adhesion, spheroid invasion, and static adhesion assays. These experiments were designed to analyse critical steps involved in the metastatic cascade. The transendothelial adhesion was evaluated by seeding KDM6A-deficient HOS cells onto a confluent EA.hy926 endothelial monolayer, followed by removal of non-adherent cells and quantification of cells adhered to the endothelial coating. Notably, loss of KDM6A significantly enhanced the adhesive capacity of HOS cells, indicative of increased colonization potential (**Fig. 2D).** Furthermore, to substantiate the enhanced invasive phenotype, a spheroid invasion assay was performed, wherein KDM6A-deficient spheroids plated on gelatin-coated surfaces exhibited pronounced radial outgrowth, quantified as a measure of collective invasion (**Fig. 2E**). Concomitantly, cell-matrix adhesion was analysed under static conditions on gelatin-coated plates, where KDM6A-deficient HOS cells showed increased adherence, reflecting elevated intrinsic adhesive potential (**Supplementary Fig 3B)**. These assays consistently demonstrated that loss of KDM6A not only increased the adhesive capacity of OS cells but also significantly promoted their invasive behaviour across the endothelial layer and the extracellular matrix.

**Figure 2.**
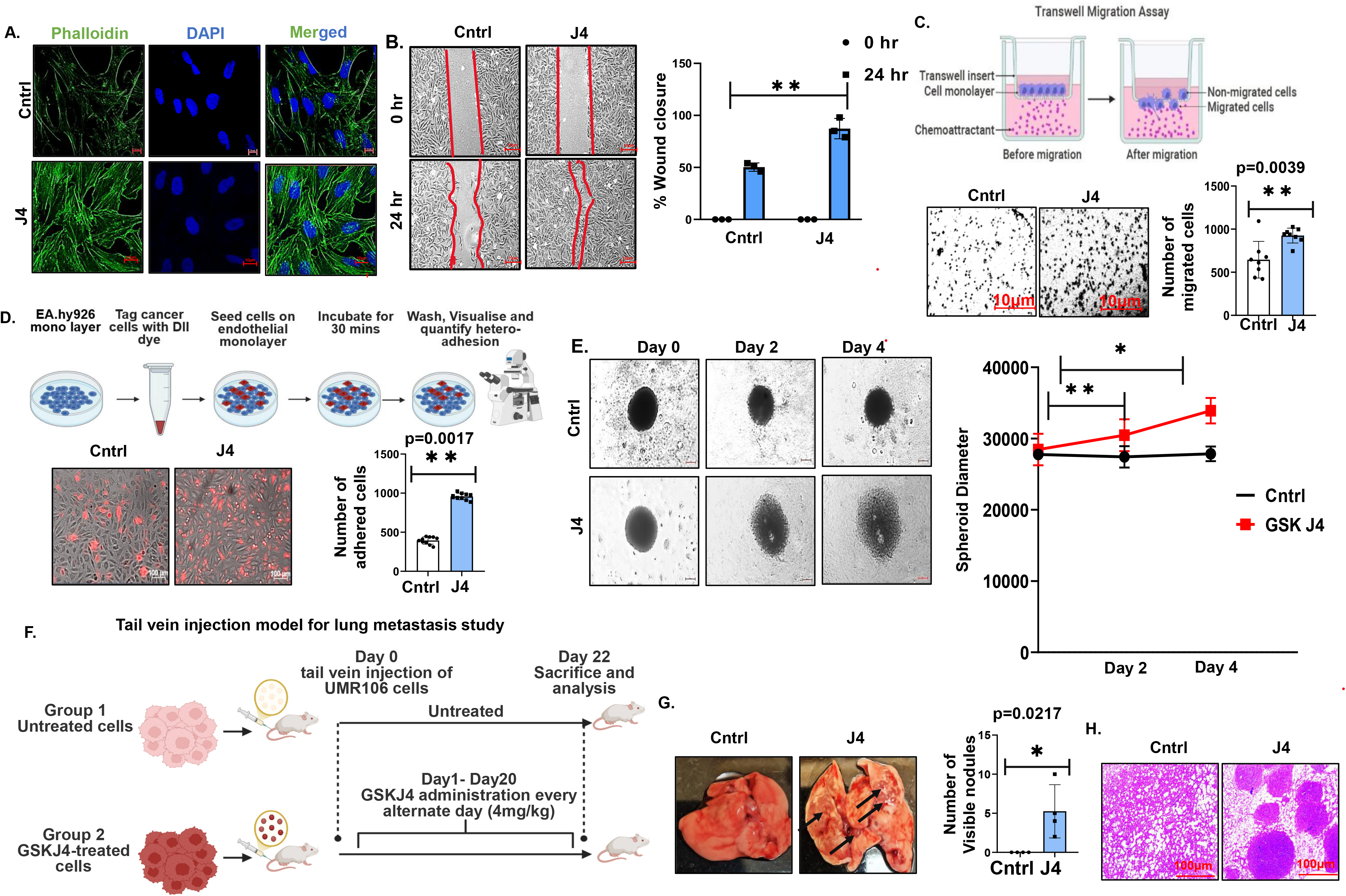
KDM6A inhibition promotes actin cytoskeleton reorganisation, cell migration, transendothelial adhesion, and spheroid growth in OS cells. **(A)** Immunofluorescence staining showing Phalloidin (F-actin) in untreated control (Cntrl) and J4-treated cells. DAPI was used to stain the nucleus. (Scale Bar: 10 μm). **(B)** Representative images of wound healing assay at 0 hr and 24 hr in J4-treated cells, and bar graph representing percentage wound closure at 24 hr in J4-treated cells (Scale Bar: 10 μm). **(C)** Representative image of transwell migration assay and quantification of the number of migrated cells in transwell migration assay in J4-treated cells (Scale Bar: 10 μm). **(D)** Representative images and quantification of the number of adhered cells in the transendothelial adhesion assay in J4-treated cells (Scale Bar: 100 μm). **(E)** Representative image showing spheroid formation at Day 0, Day 2 and Day 4 in J4-treated cells and bar graph representing quantification of spheroid diameter at different time points. **(F)** Schematic representation of the tail vein injection model of lung metastasis. J4-treated (10 μM; 24 hr) or untreated UMR106 cells were intravenously injected *via* the lateral tail vein into Sprague-Dawley rats. Additionally, animals were administered J4 (4 mg/kg) on alternate days for 20 days and sacrificed on day 22 for evaluation of pulmonary metastasis. **(G)** Representative images of lungs isolated from animals receiving KDM6A-inhibitor compared to untreated rat lungs, along with quantification of visible metastatic lung nodules. The black arrow represents a visible lung nodule in J4-treated animals. **(H)** Representative H&E-stained lung tissue sections from KDM6A-inhibited groups showing metastatic lesions (Scale Bar: 100 μm). Unless otherwise specified, treatments were conducted for 24 h, and cells were treated with the KDM6A inhibitor J4 (10 μM). All data are presented as mean ± SD from at least three independent biological replicates (n = 3). Statistical significance between two groups was determined using an unpaired two-tailed Student’s t-test. Statistical significance is indicated as (*) p<0.05, (**) p<0.005 and (***) p<0.005. Cntrl denotes untreated cells. J4, GSKJ4; DAPI, 4’,6-diamidino-2-phenylindole.

To validate whether the enhanced metastatic traits observed *in vitro* translate into increased pulmonary colonisation *in vivo*, we employed an experimental tail vein injection model of lung metastasis; a schematic representation of the study design is provided in **Fig. 2F**. UMR106 osteosarcoma cells were intravenously injected into two groups of Sprague-Dawley rats *via* the lateral tail vein, with one group receiving untreated cells and the other receiving J4-treatment. Animals were subsequently administered J4 on alternate days for 20 days and sacrificed on day 22 for evaluation of pulmonary metastasis. This model allows circulating tumour cells to colonise the lungs and recapitulates the later stages of the metastatic cascade, including vascular survival, extravasation, and pulmonary colonisation. Notably, animals injected with KDM6A-inhibited cells exhibited a significantly greater metastatic burden than the control group, as evidenced by a marked increase in the number of visible metastatic lung nodules **(Fig. 2G)**. Histopathological examination of H&E-stained lung sections further confirmed extensive metastatic lesions in the KDM6A-inhibited group **(Fig. 2H)**, demonstrating that KDM6A loss markedly promotes pulmonary metastatic colonisation *in vivo*.

An increased invasiveness is frequently associated with reduced sensitivity to anticancer therapies (29). Consistent with this notion, KDM6A inhibition–induced invasive phenotypes in OS cells were accompanied by a marked decrease in drug sensitivity. Cells treated with anticancer agent cisplatin (CDDP), which is considered one of the first-line chemotherapy drugs for OS patients, exhibited reduced apoptotic response, as evidenced by Annexin V staining (**Supplementary Fig. 3C)**; increased viability as analysed by MTT assay (**Supplementary Fig. 3D),** and enhanced survival in 3D spheroid models (**Supplementary Fig. 3E)**, following KDM6A inhibition, indicating a robust therapy resistance.

### 3.3. β-catenin expression is elevated following KDM6A inhibition in OS cells

The WNT/β-catenin signalling pathway is frequently dysregulated in OS, making it one of its important hallmarks (30). By driving EMT, modulating cell-matrix interactions, and enhancing adhesive and invasive capacities, this pathway plays a pivotal role in facilitating metastatic dissemination and therapy resistance in OS. In the previous sections, we observed increased invasiveness with KDM6A inhibition; hence, we next sought to investigate whether WNT/β-catenin signalling played a role in our context. We found that upon KDM6A inhibition, β-catenin expression increases at both the transcript **(Fig. 3A)** and protein **(Fig. 3B & 3C, Supplementary Fig. 4A)** levels, and that β-catenin expression correlates with KDM6A activity. Conversely, overexpression of KDM6A resulted in a marked reduction in β-catenin expression (**Supplementary Fig. 4B).** In addition, we also looked for the reported downstream target genes of β-catenin, such as CD44 (**Fig. 3D, Supplementary Fig. 4C)** and Cyclin D1 **(Fig. 3E)**; notably, we found out that they are also consistently upregulated upon KDM6A inhibition, further supporting the notion that loss of KDM6A activity activates the WNT/β-catenin signalling cascade. To further validate these observations, KDM6A was inhibited in another OS cell type, MG-63, where we observed comparable results **(Supplementary Fig. 4D & 4E)**. In addition, we analysed the subcellular distribution of β-catenin and CD44 using cellular fractionation and immunofluorescence assay **(Fig. 3F-G; Supplementary Fig. 4F& 4G)**. These analyses revealed that both β-catenin and CD44 levels were elevated not only in the cytoplasmic fraction but also in the nucleus. In addition, we investigated whether β-catenin transcriptionally regulates the expression of EMT-associated markers, such as Vimentin, using ChIP-qPCR. We found that β-catenin can bind to the Vimentin promoter region and regulate its expression, thereby extending its role in driving invasive phenotypes **(Fig. 3l-J)**. This adds to its previously established function, in which β-catenin was found to modulate the expression of EMT-associated transcription factors like Snail (31, 32).

**Figure 3.**
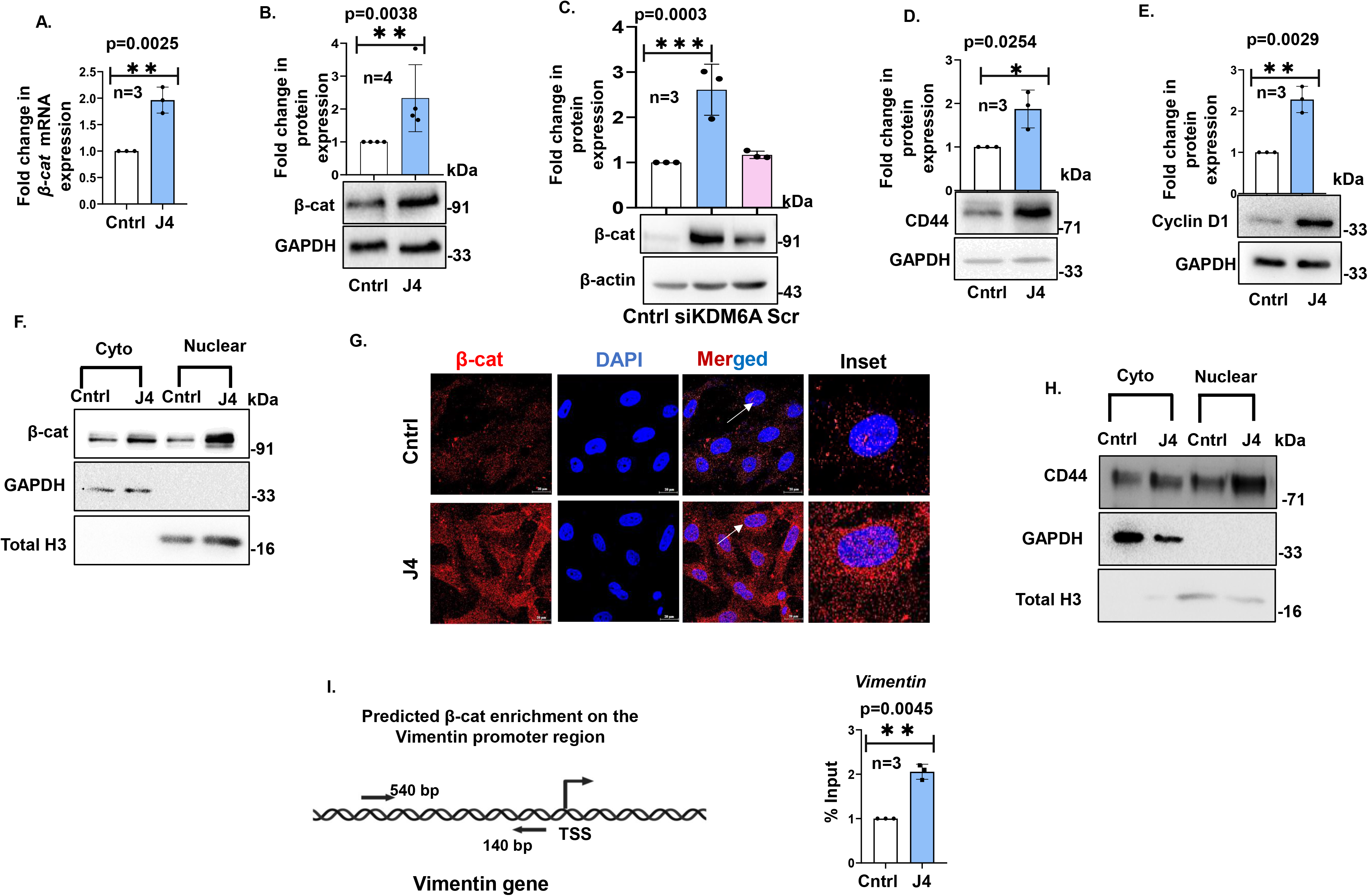
Inhibition of KDM6A upregulates β-catenin expression, promotes its nuclear translocation and enhances its recruitment to the Vimentin promoter in OS cells. **(A)** Fold change in mRNA expression of β-catenin in J4 treated cells. **(B)** Immunoblot analysis showing β-catenin expression in J4-treated cells; β-actin is the loading control. **(C)** Immunoblot analysis showing β-catenin expression upon siKDM6A transfection. Scr served as scrambled control transfected cells; β-actin is the loading control. Immunoblot analysis showing CD44 **(D)** and Cyclin D1 **(E)** expression in J4-treated cells; GAPDH is the loading control. **(F)** Immunoblot analysis showing expression of β-catenin in cytoplasmic and nuclear fractions in J4-treated cells; GAPDH and total H3 served as the loading controls for cytoplasmic and nuclear fractions, respectively. **(G)** Immunofluorescence staining showing the expression and localisation of β-catenin in J4-treated cells. Inset represents enlarged views of the regions indicated by the white arrows. (DAPI: nuclei; merged and inset images shown). **(H)** Immunoblot analysis showing CD44 expression in cytoplasmic and nuclear fractions of J4-treated cells; GAPDH and total H3 are the loading controls for the cytoplasmic and nuclear fractions, respectively. **(I)** Schematic representation of predicted β-catenin binding sites on the Vimentin promoter region, and the bar graph represents the enrichment of β-catenin over the Vimentin promoter as analysed through ChIP-qPCR in J4-treated cells. Unless otherwise specified, treatments were conducted for 24 h, and cells were treated with the KDM6A inhibitor J4 (10 μM) and siKDM6A (50 nM). Fold-change values are expressed relative to the control group, which was set to 1. All data are presented as mean ± SD from at least three independent biological replicates (n = 3). Statistical significance between two groups was determined using an unpaired two-tailed Student’s t-test. Statistical significance is indicated as (*) p<0.05, (**) p<0.005 and (***) p<0.005. Cntrl denotes untreated cells. J4, GSKJ4; β-cat, β-catenin; CD44, Cell Surface Antigen CD44; ChIP, Chromatin Immunoprecipitation; GAPDH, Glyceraldehyde 3-phosphate dehydrogenase; Scr, negative control in RNA interference (RNAi) experiments.

### 3.4. β-catenin inhibition abrogates the invasive phenotypes triggered by KDM6A inhibition

After establishing the involvement of β-catenin in KDM6A inhibition-induced invasiveness of OS cells, we further validated its role by inhibiting β-catenin using both a pharmacological inhibitor (Pyrvinium pamoate; PP) and siRNA-mediated knockdown (**Fig. 4A & Supplementary Fig. 5A)**. Notably, suppression of β-catenin under these conditions resulted in a marked reduction in the expression of N-cadherin and Vimentin at both the transcript (**Fig. 4B-C)** and protein levels (**Fig. 4D-E, Supplementary Fig. 5A & 5B)**. This was accompanied by a concomitant decrease in the expression of its downstream transcriptional targets, Cyclin D1 and CD44 **(Fig. 4F, Supplementary Fig. 5C)**. Consistently, immunofluorescence analysis also revealed a concomitant decrease in the expression of Vimentin (**Fig. 4G)** and N-cad (**Supplementary Fig. 5D)** upon β-catenin inhibition. Using transwell migration and wound-healing assays, we additionally investigated whether β-catenin inhibition could counteract the enhanced migratory and wound-healing capacities induced by KDM6A inhibition. Our results showed that blocking β-catenin led to a marked decline in wound healing **(Supplementary Fig. 6A)** and cell migration potential **(Supplementary Fig. 6B)**. We further observed that the enhanced adhesion capacity of OS cells upon KDM6A inhibition was effectively abrogated upon β-catenin inhibition (**Supplementary Fig. 6C)**. These findings indicate that inhibition of β-catenin effectively reverses KDM6A loss-induced invasiveness, thereby underscoring the pivotal role of the KDM6A–β-catenin axis in driving invasive and metastatic phenotypes in OS cells.

**Figure 4.**
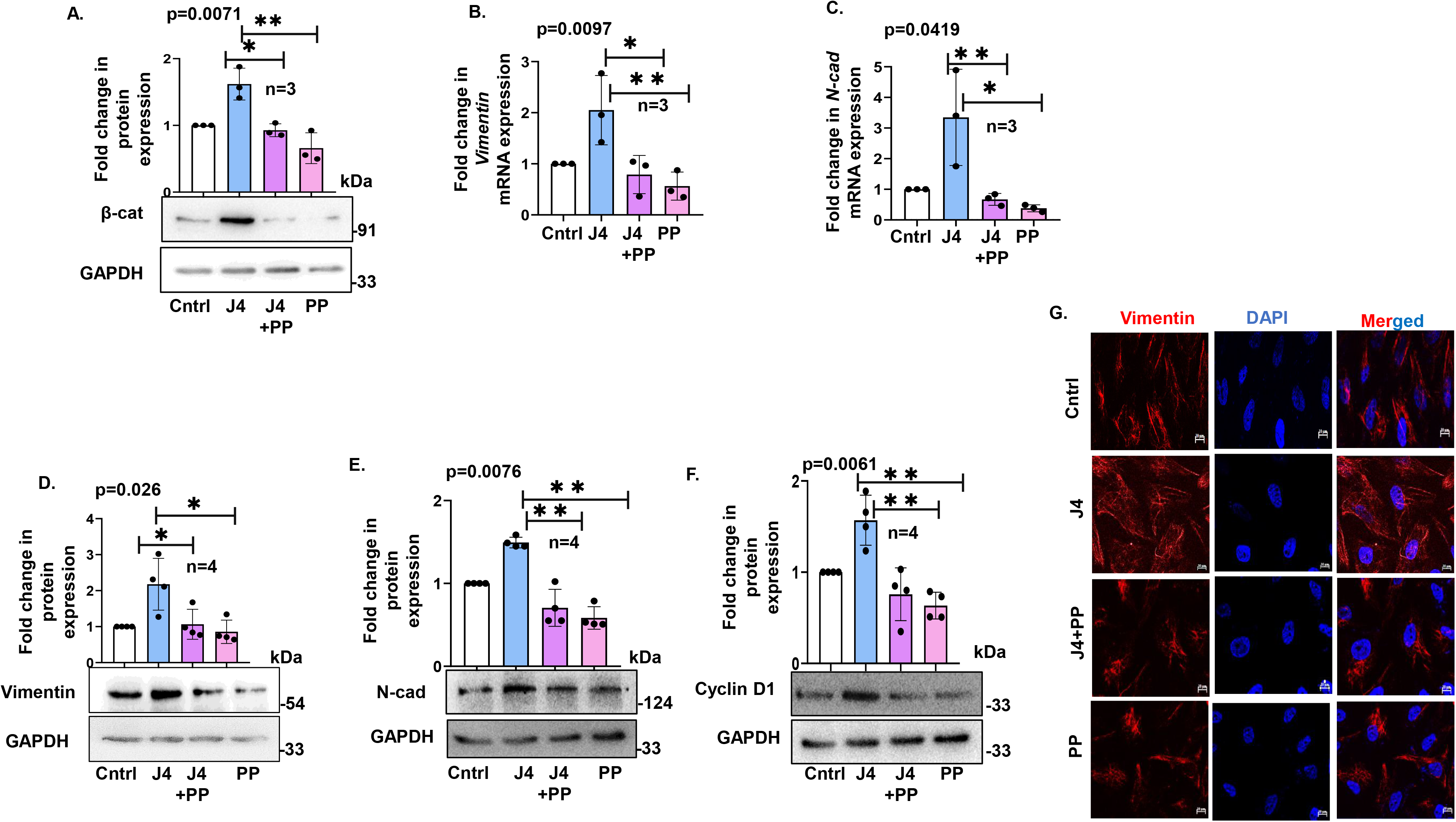
β-catenin inhibition abrogates the invasive phenotypes triggered by KDM6A inhibition. **(A)** Immunoblot analysis showing expression of β-catenin in J4, J4+PP and PP-treated cells (PP, Pyrvinium pamoate WNTi 50 nM); GAPDH is the loading control. Fold change in Vimentin **(B)** and N-cadherin **(C)** mRNA expression in J4, J4+PP and PP-treated cells. Immunoblot analysis showing expression of Vimentin **(D),** N-cadherin **(E)** and Cyclin D1 **(F)** in J4, J4+PP and PP-treated cells; GAPDH is the loading control. **(G)** Immunofluorescence staining showing the expression and localisation of Vimentin in J4, J4+PP and PP-treated cells (Scale Bar: 10 μm). Unless otherwise specified, treatments were conducted for 24 h, and cells were treated with the KDM6A inhibitor J4 (10μM) and WNTi, administered 6 hr prior to GSKJ4 treatment. Fold-change values are expressed relative to the control group, which was set to 1. All data are presented as mean ± SD from at least three independent biological replicates (n = 3). Statistical significance between two groups was determined using an unpaired two-tailed Student’s t-test. Statistical significance is indicated as (*) p<0.05, (**) p<0.005 and (***) p<0.005. Cntrl denotes untreated cells. J4, GSKJ4; β-cat, β-catenin; GAPDH, Glyceraldehyde 3-phosphate dehydrogenase; N-cad, Neural cadherin; WNTi, WNT inhibitor.

### 3.5. YAP signalling drives β-catenin expression following loss of KDM6A activity

Signalling pathways rarely operate in isolation; instead, they engage in extensive crosstalk that coordinates cellular functions. Among these, aberrant action of the YAP signalling axis and its interactions with multiple signalling pathways are often involved in tumorigenesis (21, 33). Moreover, the considerable overlap in biological functions controlled by YAP and Wnt/β-catenin suggests that they are highly interconnected and can reciprocally modulate each other’s activity (23,35,36). This bidirectional crosstalk has emerged as a critical regulatory node in cancer progression, where its dysregulation contributes to multiple steps in the tumorigenesis cascade. Therefore, in this study, we explored the potential crosstalk between YAP and Wnt/β-catenin signalling, which are often disrupted in OS (20, 21, 30). Interestingly, KDM6A inhibition led to increased H3K27me3 methylation, a closed chromatin mark at the LATS1 gene promoter (a negative regulator of YAP) **(Fig. 5A & 5B)**, which corresponded to decreased LATS1 expression at both the mRNA and protein levels **(Fig. 5C and 5D)**. Supporting these experimental findings, *in silico* analysis of the pan-cancer dataset and TARGET OS dataset from cBioPortal revealed a significant positive correlation between KDM6A expression and the upstream Hippo-signalling regulator, i.e., LATS1 **(Supplementary Fig. 7A)** and MOB1A **(Supplementary Fig. 7B)**, underlining a potential role of KDM6A loss in regulating the Hippo signalling pathway in OS. As LATS1 is an upstream kinase that phosphorylates YAP and targets it for ubiquitination-mediated degradation, we monitored YAP expression (22, 36). In accordance with the above, we observed elevated total YAP at both the transcriptional **(Fig. 5E)** and protein **(Fig. 5F)** levels, accompanied by decreased pYAPser127 expression **(Fig. 5G);** phosphorylation at this site primes the protein for degradation. This finding further supports the regulatory link between KDM6A inhibition and YAP activation. Alternatively, KDM6A overexpression reduced YAP levels **(Supplementary Fig. 7C)**. Furthermore, interestingly, cellular fractionation analysis revealed that YAP was predominantly localised in the cytoplasm rather than in the nucleus **(Fig. 5H-I)**. Consistent with this observation, canonical YAP downstream transcriptional targets like Cyr61 and CTGF did not show a trend of increase with YAP accumulation, indicating that YAP may exert non-transcriptional functions in the cytoplasm under these conditions **(Fig. 5J–M)**. To further evaluate YAP transcriptional activity, a YAP luciferase reporter assay was conducted. Notably, KDM6A inhibition didn’t alter YAP-driven reporter activity compared to untreated control (**Fig. 5N)**. This finding is particularly intriguing, as increased YAP expression is usually associated with its nuclear accumulation and subsequent activation of target gene transcription. In contrast, our data indicate that, despite its elevated expression, YAP remains largely sequestered in the cytoplasm in KDM6A-deficient cells.

**Figure 5.**
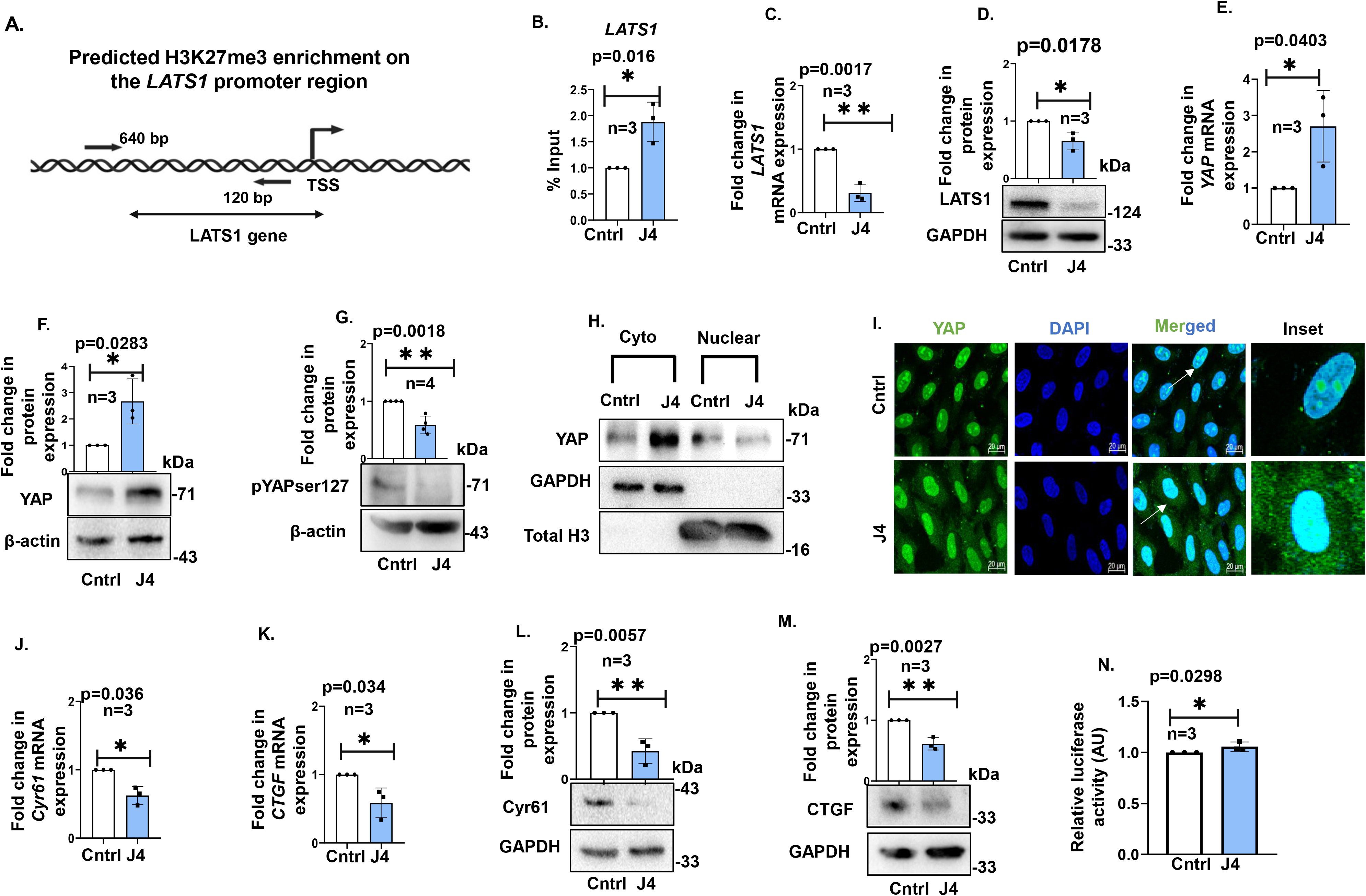
Inhibition of KDM6A downregulates LATS1 and activates YAP signalling. **(A)** Schematic representation of predicted H3K27me3 binding site on the LATS1 promoter region and ChIP-qPCR analysis showing its % Input enrichment at the LATS1 promoter in control versus J4-treated cells. **(C)** Fold change in mRNA expression of LATS1 in J4-treated cells. **(D)** Immunoblot analysis showing LATS1 expression in J4-treated cells; GAPDH is the loading control. **(E)** Fold change in mRNA expression of YAP in J4-treated cells. **(F)** Immunoblot analysis showing YAP expression J4-treated cells; β-actin is the loading control. **(G)** Immunoblot analysis showing expression of pYAP (Ser127) in J4-treated cells; β-actin is the loading control. **(H)** Immunoblot analysis showing YAP expression in cytoplasmic and nuclear fractions in J4-treated cells; GAPDH and total H3 are the loading controls. **(I)** Immunofluorescence staining showing the expression and localisation of YAP in J4-treated cells. Insets represent enlarged views of the regions indicated by the white arrows. (DAPI: nuclei; merged and inset images shown) (Scale Bar: 20 μm). Fold change in mRNA expression of Cyr61 **(J)** and CTGF **(K)** in J4-treated cells. Immunoblot analysis showing Cyr61 **(L)** and CTGF **(M)** expression in J4-treated cells; GAPDH is the loading control. **(N)** Change in luciferase activity of YAP-responsive promoter construct post KDM6A inhibition. Unless otherwise specified, treatments were conducted for 24 h, and cells were treated with the KDM6A inhibitor J4 (10 μM). Fold-change values are expressed relative to the control group, which was set to 1. All data are presented as mean ± SD from at least three independent biological replicates (n = 3). Statistical significance between two groups was determined using an unpaired two-tailed Student’s t-test. Statistical significance is indicated as (*) p<0.05, (**) p<0.005 and (***) p<0.005. Cntrl denotes untreated cells; ChIP, Chromatin Immunoprecipitation; GAPDH, Glyceraldehyde 3-phosphate dehydrogenase; LATS1, Large Tumor Suppressor kinase 1; YAP, Yes associated protein; β-actin, beta actin; Cyr61, Cysteine-rich, angiogenic inducer, 61; CTGF, Connective Tissue Growth Factor.

### 3.6. YAP regulates the expression of β-catenin in a proteasomal degradation pathway-dependent manner

Next, we investigated whether YAP influences β-catenin expression. Interestingly, *in silico* analysis of pan-cancer dataset and TARGET-OS dataset from cBioPortal revealed a significant positive correlation between YAP and β-catenin across multiple cancer types, suggesting potential crosstalk between these two evolutionarily conserved signalling pathways **(Supplementary Fig. 8A)**. To gain further mechanistic insight, YAP was inhibited either pharmacologically using VP or through siRNA and confirmed through western blot **(Fig. 6A & Supplementary Fig. 8B).** Both approaches led to a marked reduction in β-catenin protein levels **(Fig. 6B & Supplementary Fig. 8C)**, while its mRNA levels remained considered unaffected **(Fig. 6C).** This indicates that YAP regulates β-catenin abundance at the post-transcriptional level rather than through transcriptional control. Given the observed cytoplasmic accumulation of YAP, we hypothesise that YAP may influence β-catenin protein turnover through modulation of GSK3β, a key cytoplasmic kinase within the β-catenin destruction complex. Under basal conditions, in the absence of Wnt signalling, the destruction complex efficiently sequesters cytosolic β-catenin, promoting its phosphorylation by GSK3β and subsequent degradation by ubiquitin ligase (19). However, a loss of GSK3β activity, frequently associated with its inhibitory phosphorylation at Ser9, can facilitate its degradation, resulting in stabilisation and subsequent nuclear translocation of β-catenin (37). Notably, in our study KDM6A inhibition resulted in a significant reduction in GSK3β expression **(Fig. 6D, Supplementary Fig. 8D),** whereas KDM6A overexpression increased GSK3β levels **(Supplementary Fig. 8E)**. Consistent with this, we observed a marked increase in the expression of phospho-GSK3βser9 following KDM6A inhibition compared to untreated cells **(Fig. 6E)**. Furthermore, to determine whether the reduction in GSK3β was mediated through proteasomal degradation, cells were pre-treated with MG132, a proteasomal inhibitor, which restored GSK3β expression relative to KDM6A-inhibited conditions **(Supplementary Fig. 8F)**. Herein, we hypothesised that YAP forms part of the β-catenin destruction complex, as previous reports have suggested an association between YAP and components of the complex (23). Importantly, immunoprecipitation of YAP, upon KDM6A inhibition, demonstrated a robust interaction with GSK3β **(Fig. 6F)**. Consistently, co-localisation analysis further confirmed the association between YAP and GSK3β within the cytoplasm **(Fig. 6G)**. Our results point to a probable mechanistic interplay in which cytoplasmic YAP regulates β-catenin stability at the post-transcriptional level by modulating destruction complex-associated proteins. To determine this, we inhibited YAP with VP in the presence of the proteasomal inhibitor MG132. β-catenin expression was partially restored in the presence of MG132 in the KDM6A inhibition condition, indicating that proteasome-mediated degradation accounts for the reduction in β-catenin upon YAP inhibition **(Fig. 6H & Supplementary Fig. 8G)**. Furthermore, to rule out the involvement of autophagy in maintaining β-catenin expression, cells were treated with the autophagy inhibitor CQ. However, CQ treatment did not produce any significant change in β-catenin levels, suggesting that autophagy does not contribute substantially to β-catenin maintenance in this context (**Supplementary Fig. 8H**). To directly assess whether β-catenin stability was linked to altered ubiquitination, we performed β-catenin pull-down assays and probed for ubiquitin. Interestingly, we observed reduced ubiquitin conjugation to β-catenin under KDM6A-inhibited conditions, suggesting that YAP protect β-catenin from ubiquitin-mediated proteasomal degradation **(Fig. 6I)**. Taken together, these results provide strong evidence that YAP interacts with the β-catenin destruction complex and contributes to the stabilisation of β-catenin protein in this context.

**Figure 6.**
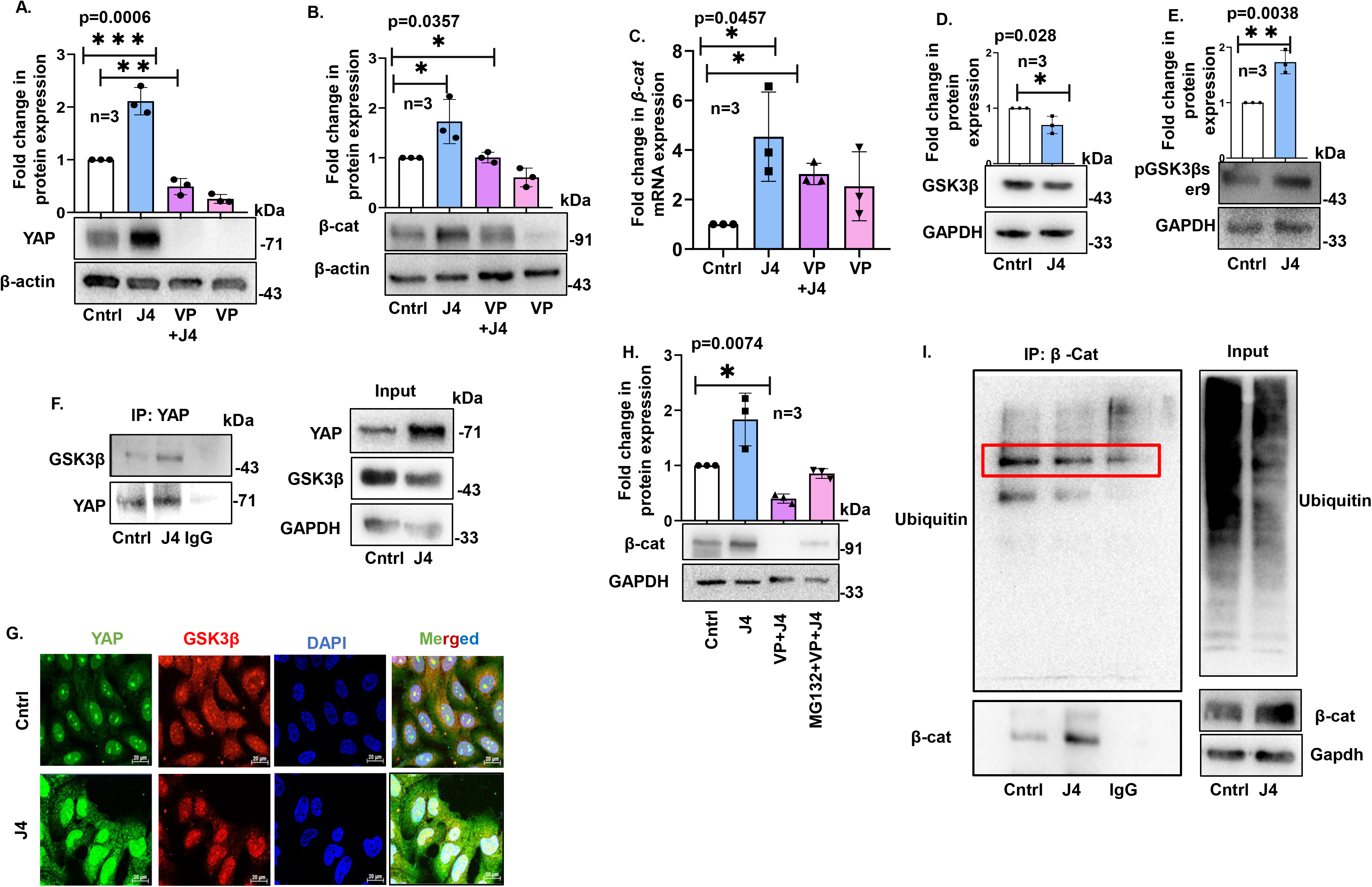
KDM6A inhibition by J4 inactivates GSK3β, promotes YAP-GSK3β interaction, and stabilises β-catenin by reducing its ubiquitination in OS cells. **(A)** Immunoblot analysis showing expression of YAP and β-catenin **(B)** in J4, J4+VP, and VP-treated cells (VP, YAPi, 5 μM); β-actin is the loading control. **(C)** Fold change in mRNA expression of β-catenin in J4, J4+VP, and VP-treated cells. Immunoblot analysis showing GSK3β **(D)** and pGSK3βser9 **(E)** expression in J4-treated cells; GAPDH is the loading control. **(F)** Co-immunoprecipitation analysis (IP: YAP) showing interaction between YAP and GSK3β in J4-treated cells; Input and GAPDH are shown. **(G)** Immunofluorescence staining showing co-localisation of YAP and GSK3β in J4-treated cells (DAPI: nuclei; merged images shown) (Scale Bar: 20 μm). **(H)** Immunoblot analysis showing expression of β-catenin in J4, VP+J4, MG132+VP+J4, VP and MG132 treated cells (MG132, proteasome inhibitor, 0.5 μM); GAPDH is the loading control. **(I)** Co-immunoprecipitation analysis (IP: β-catenin) followed by immunoblot for ubiquitin in J4-treated cells; IgG control shown. Unless otherwise specified, treatments were conducted for 24 h, and cells were treated with the KDM6A inhibitor J4 (10μM) or VP or MG132. If only VP is treated prior to GSKJ4, it is treated for 6 hr; if both MG132 and VP are treated together, each is treated for 3 hr prior to GSKJ4. Fold-change values are expressed relative to the control group, which was set to 1. All data are presented as mean ± SD from at least three independent biological replicates (n = 3). Statistical significance between two groups was determined using an unpaired two-tailed Student’s t-test. Statistical significance is indicated as (*) p<0.05, (**) p<0.005 and (***) p<0.005. Cntrl denotes untreated cells. J4, GSKJ4; VP, Verteporfin; DAPI, 4’,6-diamidino-2-phenylindole; β-cat, β-catenin; GAPDH, Glyceraldehyde 3-phosphate dehydrogenase; GSk3β, Glycogen Synthase Kinase-3 Beta; YAP, Yes Associated Protein; pGSK3βser9, Phospho-Glycogen Synthase Kinase-3 Beta serine 9; YAPi, YAP inhibitor.

## 4. Discussion

Metastasis remains the primary cause of mortality in OS, a highly aggressive primary bone malignancy that predominantly affects children and adolescents. Despite aggressive multimodal therapy, patients with metastatic disease continue to face dismal outcomes, with five-year survival rates falling below 20% (5). This poor prognosis stems largely from the tumour’s strong propensity for early hematogenous spread, most commonly to the lungs. While extreme genomic instability is a well-recognised hallmark of OS, emerging evidence highlights epigenetic dysregulation, particularly involving histone modifiers, as a critical driver of its metastatic behaviour (38).

The present study demonstrates that loss of the histone demethylase KDM6A drives a pronounced mesenchymal and invasive phenotype in OS cells through coordinated activation of Wnt/β-catenin signalling and non-canonical modulation of the Hippo/YAP axis. We show that KDM6A deficiency markedly enhances migratory, adhesive, and invasive capacities, as evidenced by upregulation of mesenchymal markers and downregulation of epithelial markers. While KDM6A mutations or downregulation have been extensively documented across multiple cancer types and are consistently associated with reduced patient survival and increased metastatic risk, their functional role in OS has remained largely unexplored. Previous clinical studies in bladder, breast, myeloid leukaemia and adenoid cystic carcinomas have shown that low KDM6A expression or loss-of-function mutations correlate with higher metastatic rates and poorer outcomes (15, 16, 39, 40, 41). Our *in-silico* analysis of OS datasets now extends these observations by revealing a strong association between KDM6A deficiency and lung metastasis in OS — a finding that aligns with the dismal prognosis of metastatic OS. Thus, our work positions KDM6A loss as a clinically relevant driver of metastatic behaviour, specifically in this aggressive bone malignancy.

Furthermore, we observed that KDM6A loss resulted in upregulation of mesenchymal markers (Vimentin, N-cadherin) and downregulation of epithelial markers (E-cadherin). Importantly, this activation also enhanced trans-endothelial adhesion — a critical step that enables circulating tumour cells to adhere to the vascular endothelium and extravasate during metastatic dissemination — accompanied by enhanced cytoskeletal remodelling and induced migration (42). These phenotypic changes were further linked to reduced cisplatin sensitivity. We also identified a direct association of KDM6A with the Vimentin promoter, providing a mechanistic basis for its role. Both pharmacological and genetic inhibition of KDM6A induced the invasive phenotype, whereas KDM6A overexpression reversed these effects, establishing a causal link between KDM6A deficiency and heightened invasiveness in OS.

A central finding of this study is also the activation of Wnt/β-catenin signalling upon suppression of KDM6A. Although dysregulation of the Wnt/β-catenin axis is well recognised in OS and linked to advanced tumour stage and metastasis, the upstream epigenetic control by KDM6A has not been described earlier. We show that KDM6A loss leads to β-catenin accumulation, decreased GSK3β levels, and upregulation of downstream transcriptional targets (cyclin D1, CD44). Pharmacological inhibition of β-catenin completely reversed the KDM6A-loss-induced migration, adhesion, and invasion, confirming that β-catenin is a key mediator of these effects. Our study further uncovered a non-canonical mechanism linking KDM6A to the Hippo pathway. As KDM6A functions as a specific H3K27 demethylase, its inhibition leads to increased deposition of repressive H3K27me3 marks on the LATS1 promoter. This epigenetic silencing downregulates LATS1 expression, thereby reducing phosphorylation of YAP at Ser127. Interestingly, rather than undergoing canonical nuclear translocation, YAP accumulated predominantly in the cytoplasm under KDM6A-deficient conditions. This cytoplasmic YAP was found to be required for β-catenin stability by preventing its ubiquitin-mediated proteasomal degradation, thereby amplifying Wnt/β-catenin signaling and driving invasion. Taken together, these results support a model in which KDM6A acts as an epigenetic controller that maintains LATS1 expression, restrains Wnt/β-catenin activity, and prevents metastatic progression. Loss of KDM6A disrupts this regulatory network *via* repressive histone methylation at the upstream genetic element of LATS1, leading to cytoplasmic accumulation of YAP, β-catenin stabilisation, acquisition of enhanced mesenchymal features, and ultimately an enhanced invasive phenotype in OS (**Fig. 7**).

**Figure 7.**
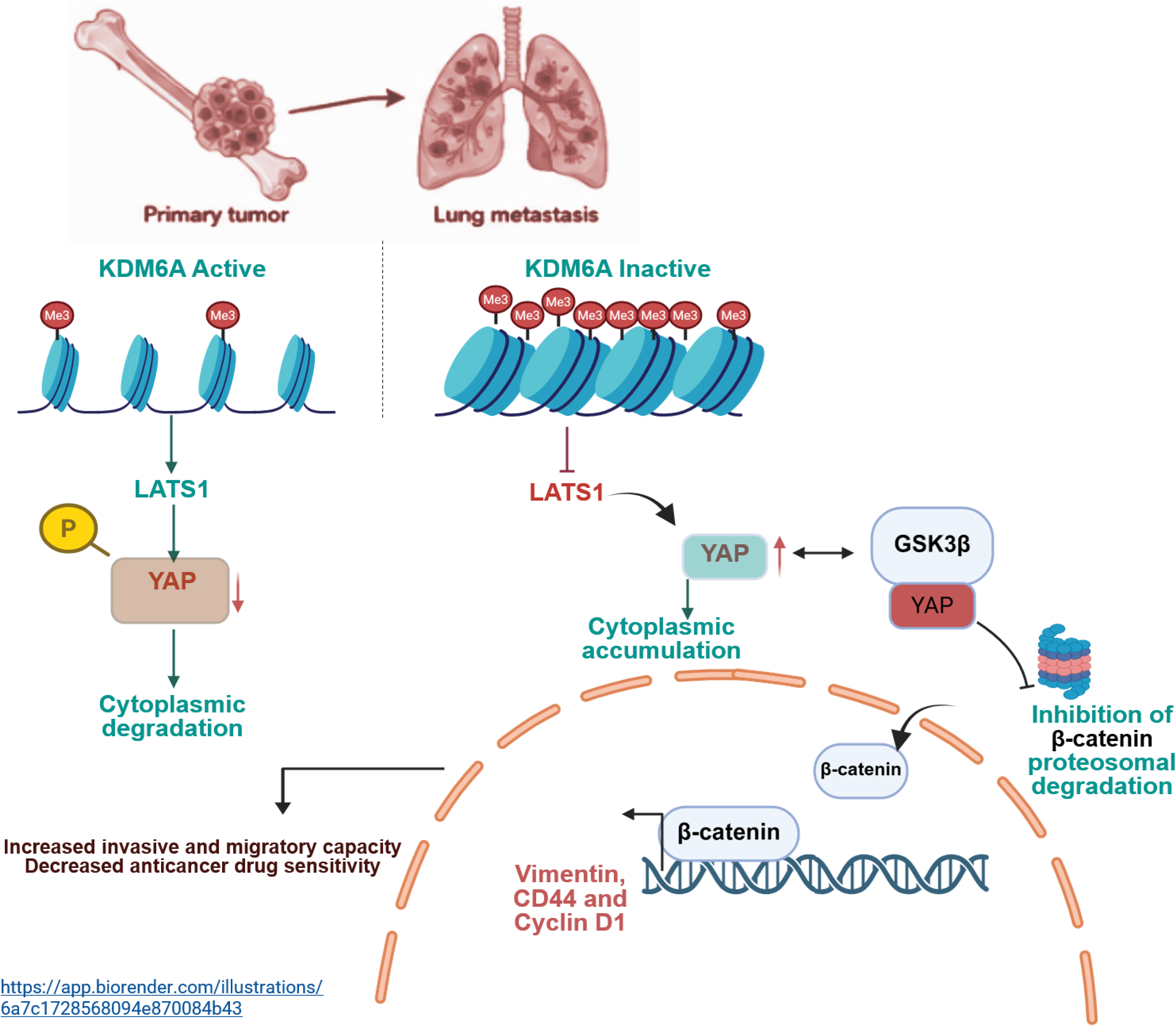
A Schematic diagram representing the KDM6A–LATS1–YAP–β-catenin axis in osteosarcoma metastasis.

From a therapeutic perspective, these findings suggest that targeting the β-catenin/YAP axis may counteract the pro-metastatic consequences of KDM6A deficiency. Pharmacological inhibitors such as pyrvinium pamoate (β-catenin) or verteporfin (YAP) effectively suppressed mesenchymal traits and invasion in KDM6A-deficient cells, highlighting a potential strategy for treating aggressive, KDM6A-deficient OS. In conclusion, this study identifies KDM6A as a key regulator of invasion and metastasis in OS and uncovers a novel epigenetic mechanism that links KDM6A-mediated regulation of LATS1 to β-catenin/YAP crosstalk.

## Authorship contribution statement

CN: Writing – original draft, Investigation, Methodology, Formal analysis, Visualisation, Data curation. MS: Animal experiment. SC: Supervision, Funding acquisition, Writing – review & editing. SM: Conceptualisation, Supervision, Funding acquisition, Writing – review & editing. RC (Corresponding Author): Conceptualisation, Supervision, Project administration, Funding acquisition, Writing – original draft, review & editing.

## Acknowledgments

CN gratefully acknowledges BITS Pilani for providing fellowship. MS gratefully acknowledges the K R Shroff Foundation for the fellowship. This research work was funded by three projects of the Department of Science and Technology, Government of India, New Delhi, under the SERB-ANRF (CRG/2022/003241), DBT (BT/PR53025/MED/30/2546/2024 dated 01.5.2025) and CDRF Project (47-06/03/8046). The authors are grateful to the Birla Institute of Technology and Science (BITS) Pilani, Pilani Campus, for providing lab facilities and infrastructure.

## Data Availability

All data supporting the findings of this study are available within the article and its Supplementary Information files or from the corresponding author upon reasonable request.

---

**Supplementary Figure 1. (A)** Relative KDM6A mRNA expression in untreated HOS (OS) and drug-tolerant persister (OS-P) HOS cells. **(B)** Pan-cancer analysis showing the number of samples with shallow deletion, diploid, gain, and amplification as obtained from cBioportal. **(C)** Kaplan–Meier analysis of overall survival of cancer patients in the pan-cancer cohort, stratified by KDM6A alteration status, as obtained from cBioportal. **(D)** Immunoblot analysis showing H3K27me3 levels in KDM6A-silenced (siKDM6A) HOS cells, with total H3 used as a loading control. (**E)** Immunoblot analysis showing H3K27me3 levels in deferiprone (DFP)-treated HOS cells. Total H3 used as loading control. **(F)** Immunoblot analysis showing the expression of KDM6A, Vimentin and N-cadherin in KDM6A-silenced (siKDM6A) HOS cells. GAPDH was used as a loading control. **(G**) Immunoblot analysis showing the expression of N-cadherin, E-cadherin, and Vimentin in DFP-treated HOS cells. GAPDH is used as a loading control. Fold-change values are expressed relative to the control group, which was set to 1. All data are presented as mean ± SD from at least three independent biological replicates (n = 3). Statistical significance between two groups was determined using an unpaired two-tailed Student’s t-test. Statistical significance is indicated as (*) p<0.05, (**) p<0.005 and (***) p<0.005. Cntrl denotes untreated cells; Scr, scrambled siRNA (Negative control); DFP, deferiprone; J4, GSKJ4, KDM6A inhibitor; KDM6A, lysine demethylase 6A; H3K27me3, trimethylated histone H3 lysine 27; H3, histone H3; siKDM6A, KDM6A-specific siRNA.

**Supplementary Figure 2. (A)** Immunoblot analysis showing the expression of N-cadherin and Vimentin **(B)** in J4-treated MG-63 cells; GAPDH used as loading control. **(C)** Immunoblot analysis showing the expression of E-cadherin and **(D)** N-cadherin in J4-treated UMR106 cells, GAPDH is used as a loading control. **(E)** Immunoblot analysis showing KDM6A and **(F)** HA-tagged KDM6A expression in KDM6A-overexpressing (OE KDM6A, KDM6A OE vector, 2 μg/ml) HOS cells, GAPDH used as loading control. **(G)** Immunoblot analysis showing H3K27me3 expression in KDM6A-OE HOS cells; GAPDH was used as a loading control, normalised to total H3. **(H)** Immunoblot analysis showing N-cadherin and **(I)** Vimentin expression in KDM6A-overexpressing HOS cells; GAPDH used as loading control. **(J)** Immunoblot analysis showing KDM6A and HA-tagged KDM6A expression **(K)** in KDM6A-OE MG-63 cells; GAPDH was used as a loading control. **(L)** Immunoblot analysis showing E-cadherin and Vimentin **(M)** expression in KDM6A-OE MG-63 cells, GAPDH was used as loading control. Fold-change values are expressed relative to the control group, which was set to 1. All data are presented as mean ± SD from at least three independent biological replicates (n = 3). Statistical significance between two groups was determined using an unpaired two-tailed Student’s t-test. Statistical significance is indicated as (*) p<0.05, (**) p<0.005 and (***) p<0.005. Cntrl denotes untreated cells; J4, GSKJ4, KDM6A inhibitor; KDM6A, lysine demethylase 6A; KDM6A-OE-KDM6A overexpression

**Supplementary Figure 3. (A)** Representative image and quantification showing percentage wound closure of wound healing assay at 0 hr, 24 hr, and 48 hr KDM6A-overexpressing (KDM6A OE vector, 2 μg/ml) cells (Scale Bar: 100 μm). **(B)** Representative image of adhered cells and its quantification in J4-treated condition (Scale Bar: 100 μm). **(C)** Quantification of Annexin V-positive cells in cisplatin (CDDP)-treated, GSKJ4 plus CDDP-treated, and J4-treated cells (CDDP; 12 μM) **(D)**. Quantification of cell viability in CDDP-treated, J4 plus CDDP-treated cells and J4-treated cells. **(E)** Representative images of spheroid formation at Day 0, Day 4, and Day 6 in CDDP-treated, DFP plus CDDP-treated, and DFP-treated cells, along with quantification of spheroid diameter at different time points. Unless otherwise specified, cells were treated with J4 and/or CDDP for 24 h. All data are presented as mean ± SD from at least three independent biological replicates (n = 3). Statistical significance between two groups was determined using an unpaired two-tailed Student’s t-test. Statistical significance is indicated as (\**) p<0.05, (**) p<0.005 and **(\*\*\****) p<0.005. Cntrl denotes untreated cells; J4, GSK J4 DFP; Deferiprone, CDDP, cis-diamminedichloroplatinum (II); KDM6A OE, KDM6A overexpression; Annexin V, Annexin V-positive apoptotic cells.

**Supplementary Figure 4. (A)** Immunoblot analysis showing β-catenin expression in DFP-treated HOS cells (DFP; 60 μM); GAPDH was used as a loading control. **(B)** Immunoblot analysis showing β-catenin expression in KDM6A-overexpressing (OE KDM6A; KDM6A OE vector, 2 μg/ml) HOS cells; GAPDH was used as a loading control. **(C)** Immunoblot analysis showing CD44 expression in DFP-treated HOS cells; GAPDH was used as a loading control. **(D)** Immunoblot analysis showing the expression of β-catenin, CD44, and CyclinD1 **(E)** in J4-treated MG-63 cells; GAPDH was used as a loading control. **(F)** Immunoblot analysis of β-catenin expression in cytoplasmic and nuclear fractions of J4-treated HOS cells. GAPDH and total H3 were used as cytoplasmic and nuclear fractionation controls, respectively. **(G)** Immunofluorescence staining showing β-catenin localisation and expression in DFP-treated HOS cells. DAPI was used to stain the nucleus. (Scale Bar: 10 μm). Unless otherwise specified, cells were treated with DFP, GSKJ4 or KDM6A OE vector for 24 h. Fold-change values are expressed relative to the control group, which was set to 1. All data are presented as mean ± SD from at least three independent biological replicates (n = 3). Statistical significance between two groups was determined using an unpaired two-tailed Student’s t-test. Statistical significance is indicated as (\**) p<0.05, (**) p<0.005 and **(\*\*\****) p<0.005. Cntrl denotes untreated cells; DFP, deferiprone; J4, GSKJ4; KDM6A OE, KDM6A overexpression; β-cat, β-catenin; CD44, cluster of differentiation 44.

**Supplementary Figure 5. (A)** Immunoblot analysis showing the expression of β-catenin and N-cadherin in J4, J4 plus β-catenin siRNA (siβ-cat)-treated, siβ-cat and Scr control (siβ-cat; 60 nM); β-actin is the loading control. **(B)** Immunoblot analysis showing Vimentin expression in J4, J4 plus siβ-Cat, siβ-Cat and Scr control (siβ-catenin, 60 nM); β-actin is the loading control. **(C)** Immunoblot analysis showing CD44 expression in DFP, DFP plus PP, and PP-treated HOS cells (DFP, 60 μM; PP, 50 nM); β-actin is the loading control. **(D)** Immunofluorescence staining showing N-cadherin expression in DFP, DFP plus PP, and PP-treated HOS cells. DAPI was used to stain the nucleus (Scale Bar: 10 μm). Unless otherwise specified, treatments were conducted for 24 h, and cells were treated with the KDM6A inhibitor J4 (10 μM), DFP, WNTi (PP) and siβ-cat. Where indicated, siβ-cat or WNT inhibitor (WNTi) treatment was administered 6 h before J4 or DFP treatment, Scr, scrambled siRNA (Negative control). Fold-change values are expressed relative to the control group, which was set to 1. All data are presented as mean ± SD from at least three independent biological replicates (n = 3). All data are presented as mean ± SD from at least three independent biological replicates (n = 3). Statistical significance between two groups was determined using an unpaired two-tailed Student’s t-test. Statistical significance is indicated as (*\*) p<0.05, (**) p<0.005 and (\*\*\**) p<0.005. Cntrl denotes untreated cells; DFP, deferiprone; J4, GSKJ4; PP, pyrvinium pamoate; siβ-cat, β-catenin-specific siRNA; β-cat, β-catenin; N-Cad, N-cadherin.

**Supplementary Figure 6. (A)** Representative images and quantification of wound healing assay at 0 hr and 24 hr in DFP, DFP plus Pyrvinium pamoate (PP), and PP-treated HOS cells (Scale Bar: 100 μm) (DFP, 60 μM; PP, 50 nM). **(B)** Representative images of transwell-migrated cells and their quantification in DFP, DFP plus PP, and PP-treated HOS cells (Scale Bar: 100 μm). **(C)** Representative images and quantification of the number of adhered cells in the static adhesion assay in DFP-treated, DFP plus PP-treated, and PP-treated HOS cells. Unless otherwise specified, treatments were conducted for 24 h, and cells were treated with the KDM6A inhibitor J4 (10 μM), DFP, and WNTi (PP). Where indicated, WNT inhibitor (WNTi) treatment was administered 6 h prior to J4 or DFP treatment. All data are presented as mean ± SD from at least three independent biological replicates (n = 3). All data are presented as mean ± SD from at least three independent biological replicates (n = 3). Statistical significance between two groups was determined using an unpaired two-tailed Student’s t-test. Statistical significance is indicated as (*\*) p<0.05, (**) p<0.005 and (\*\*\**) p<0.005. Cntrl denotes untreated cells; DFP, deferiprone; J4, GSKJ4; PP, pyrvinium pamoate; N-Cad

**Supplementary Figure 7. (A)** Correlation analysis showing the relationship between KDM6A, LATS1 and MOB1 **(B)** expression in the pan-cancer cohort as obtained from cBioportal **(C)**. Immunoblot analysis showing YAP expression in KDM6A-overexpressing (OE) HOS cells (KDM6A OE vector, 2 μg/ml); GAPDH was used as a loading control. All data are presented as mean ± SD from at least three independent biological replicates (n = 3). Fold-change values are expressed relative to the control group, which was set to 1. Statistical significance between two groups was determined using an unpaired two-tailed Student’s t-test. Statistical significance is indicated as (*\*) p<0.05, (**) p<0.005 and (\*\*\**) p<0.005. Cntrl denotes untreated cells; OE, KDM6A-overexpressing cells; KDM6A, lysine demethylase 6A; LATS1, large tumor suppressor kinase 1; YAP, Yes-associated protein; YAP1, Yes-associated protein 1.

**Supplementary Figure 8. (A)** Correlation analysis showing the relationship between YAP1 and CTNNB1 expression in the pan-cancer cohort as obtained from cBioportal. (**B**) Immunoblot analysis showing YAP and β-catenin **(C)** expression in DFP, DFP plus YAP-specific siRNA (siYAP), siYAP, and scrambled control (Scr)-treated HOS cells (DFP, Deferiprone; 50 μM; siYAP, 50 nM). GAPDH is the loading control. **(D)** Immunoblot analysis showing GSK3β expression in DFP-treated HOS cells, GAPDH is the loading control. **(E)** Immunoblot analysis showing GSK3β expression in KDM6A-overexpressing HOS cells, GAPDH is the loading control. **(F)** Immunoblot analysis showing GSK3β expression in J4, J4 plus MG132 (proteasome inhibitor, 0.5 μM), and MG132-treated HOS cells; GAPDH is the loading control. **(G)** Immunofluorescence staining showing β-catenin expression and localisation in J4, VP plus J4, and MG132 plus VP plus J4-treated HOS cells. DAPI was used to stain the nucleus. (VP, YAPi-5 μM) (Scale Bar: 10 μm). **(H)** Immunoblot analysis showing β-catenin expression in control, J4, J4 plus VP, CQ plus VP plus J4-treated, HOS cells (CQ; 10μM); GAPDH is the loading control. Unless otherwise specified, treatments were conducted for 24 h, and cells were treated with the KDM6A inhibitor J4 (10μM) or DFP or VP or MG132 or siRNA. If only VP or siRNA is treated prior to J4, it is treated for 6 hr; if both CQ and VP are treated together, each is treated for 3 hr prior to J4. Fold-change values are expressed relative to the control group, which was set to 1. All data are presented as mean ± SD from at least three independent biological replicates (n = 3). Statistical significance between two groups was determined using an unpaired two-tailed Student’s t-test. Statistical significance is indicated as (*\*) p<0.05, (**) p<0.005 and (\*\*\**) p<0.005. Cntrl denotes untreated cells; DFP, deferiprone; J4, GSKJ4; siYAP, YAP-specific siRNA; Scr, scrambled siRNA (Negative control); YAP, Yes-associated protein; β-cat, β-catenin; GSK3β, glycogen synthase kinase 3 beta; MG132, proteasome inhibitor; VP, verteporfin; CQ, chloroquine; KDM6A OE, KDM6A-overexpressing cells; CTNNB1, catenin beta 1; YAP1, Yes-associated protein 1.

