## Supplementary Figures for "KDM6A Loss Confers an Invasive Phenotype in Osteosarcoma by Activating Cytoplasmic YAP-Dependent β-catenin Stabilisation"

Supplementary Figure 1

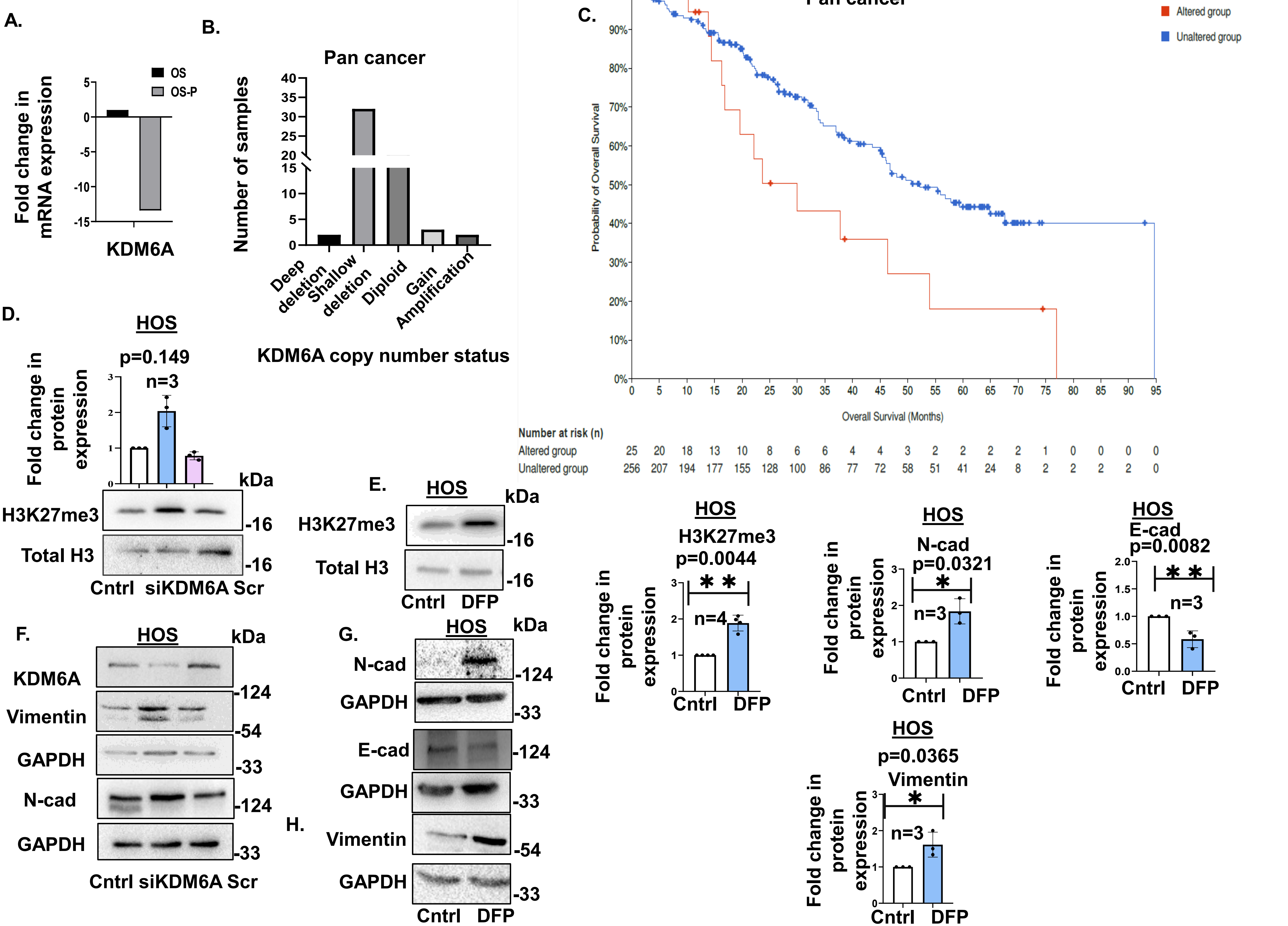

Supplementary Figure 2

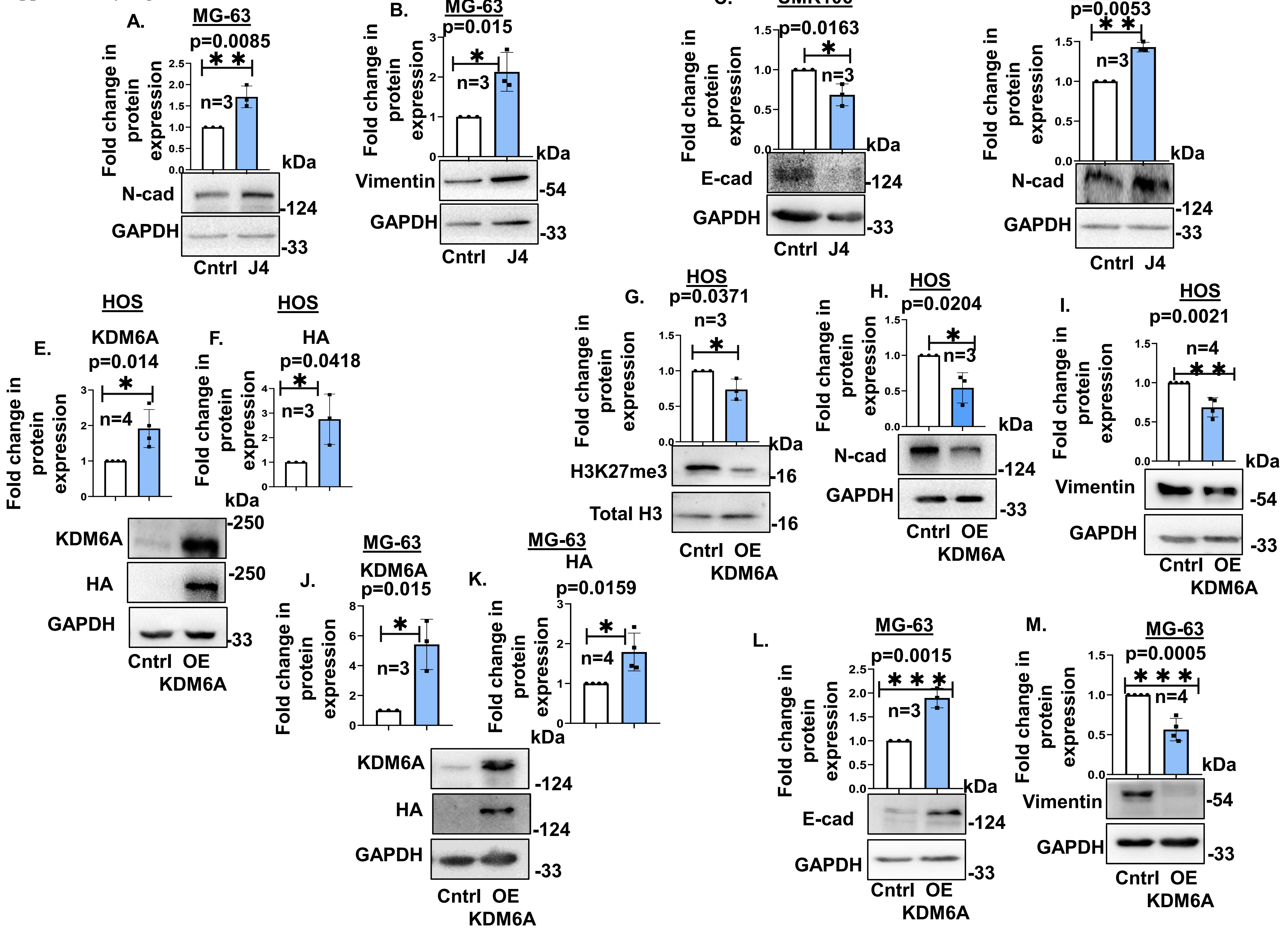

Supplementary Figure 3

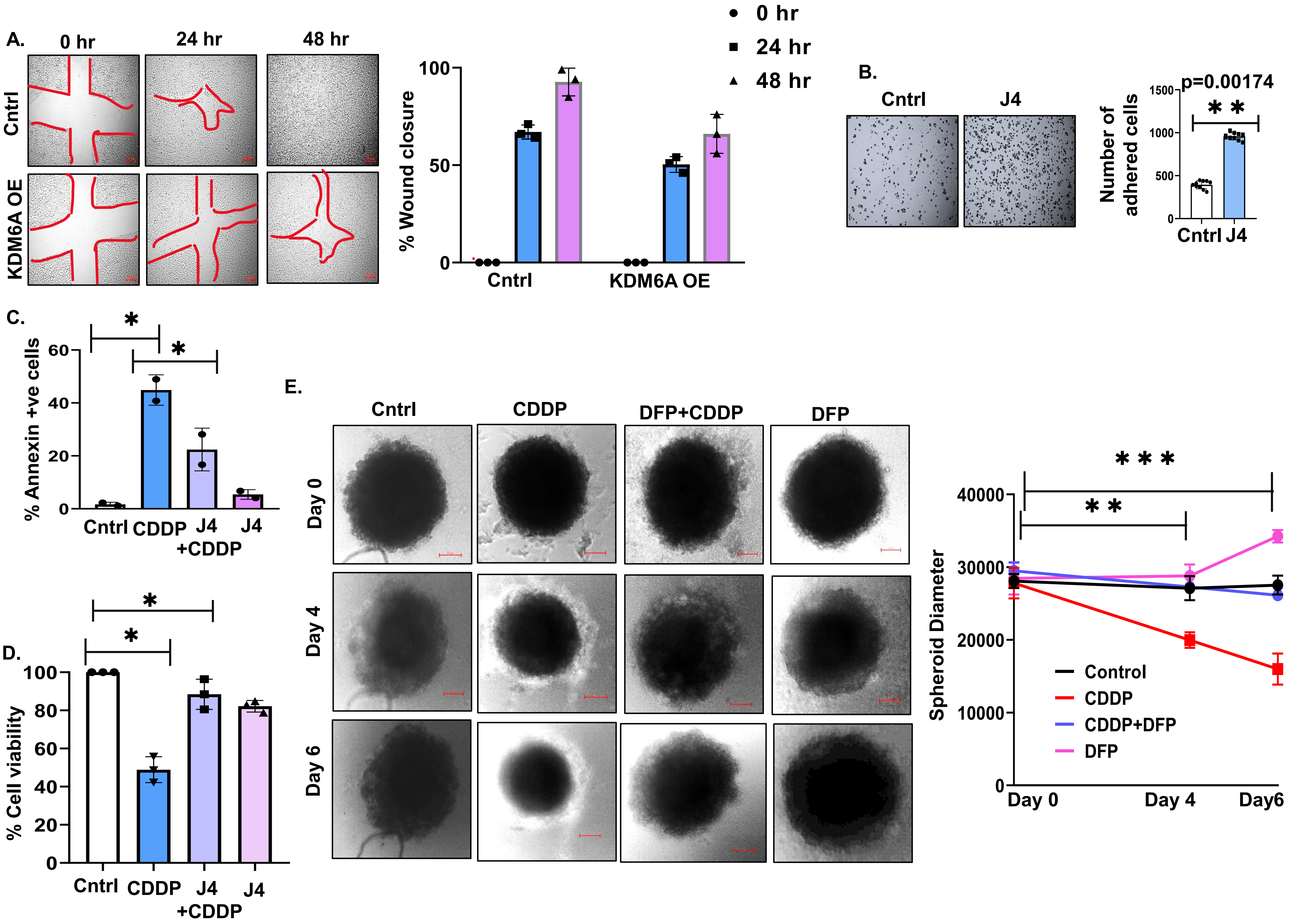

Supplementary Figure 4

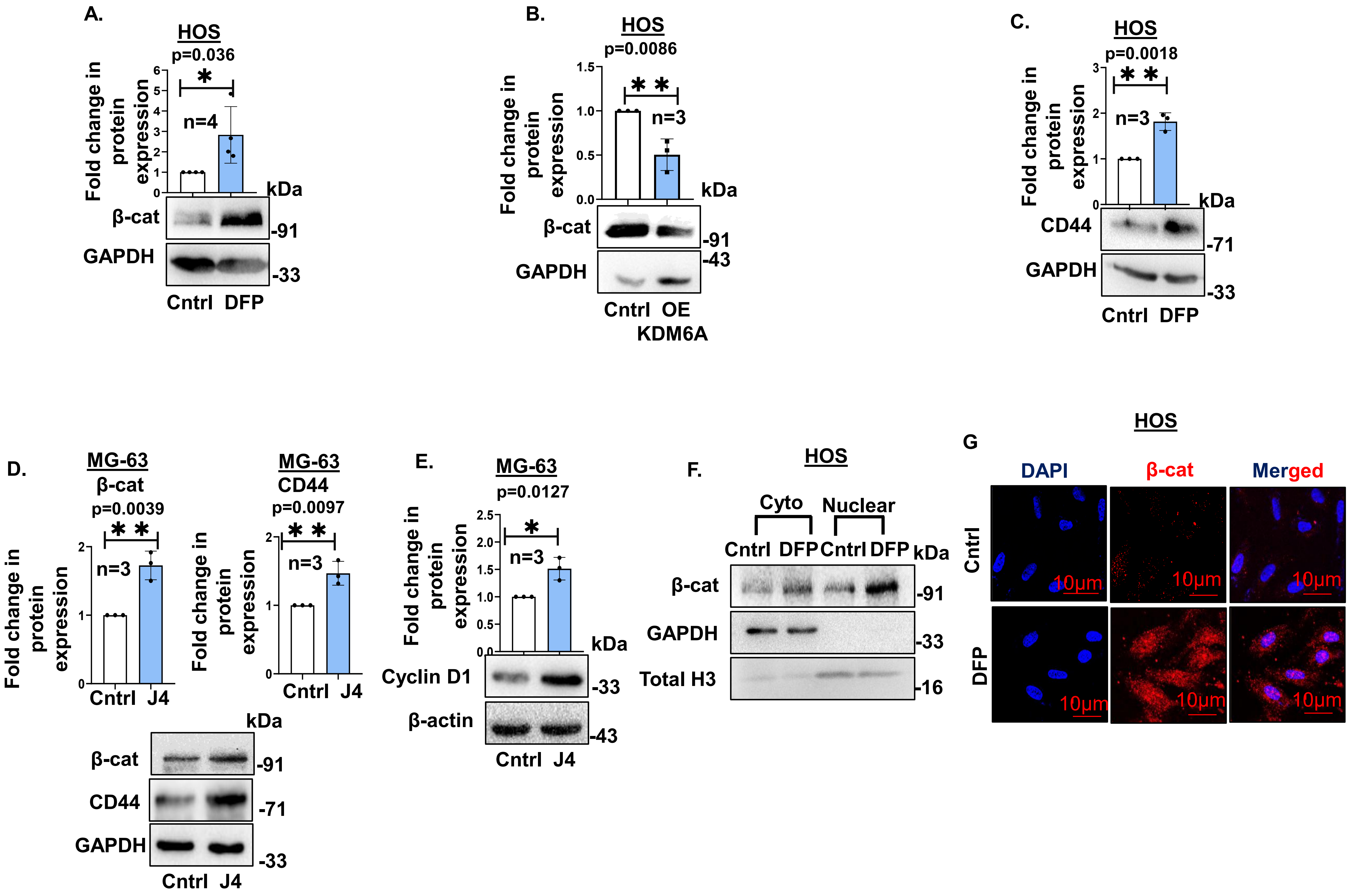

**Supplementary Figure 5**

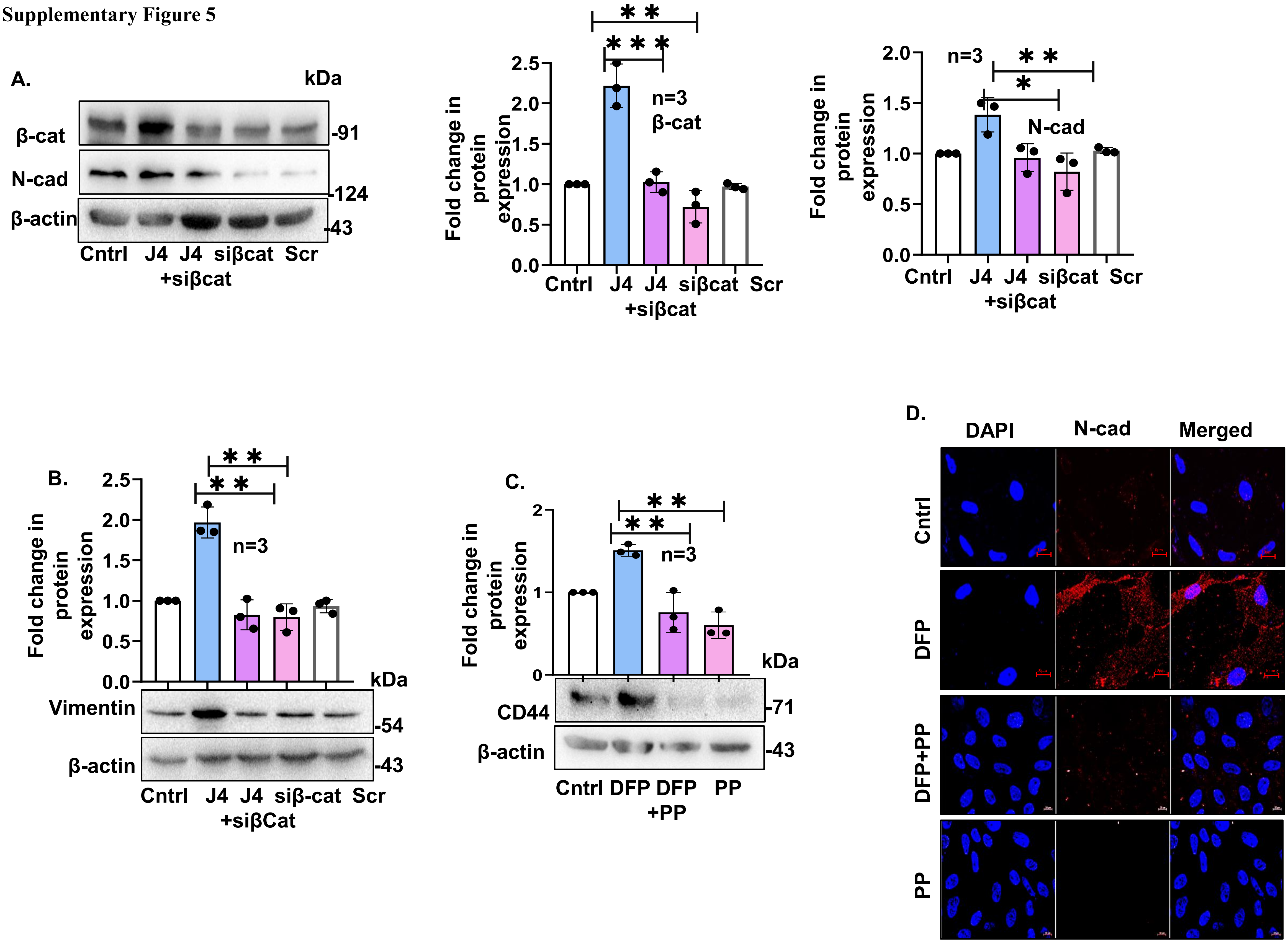

Supplementary Figure 6

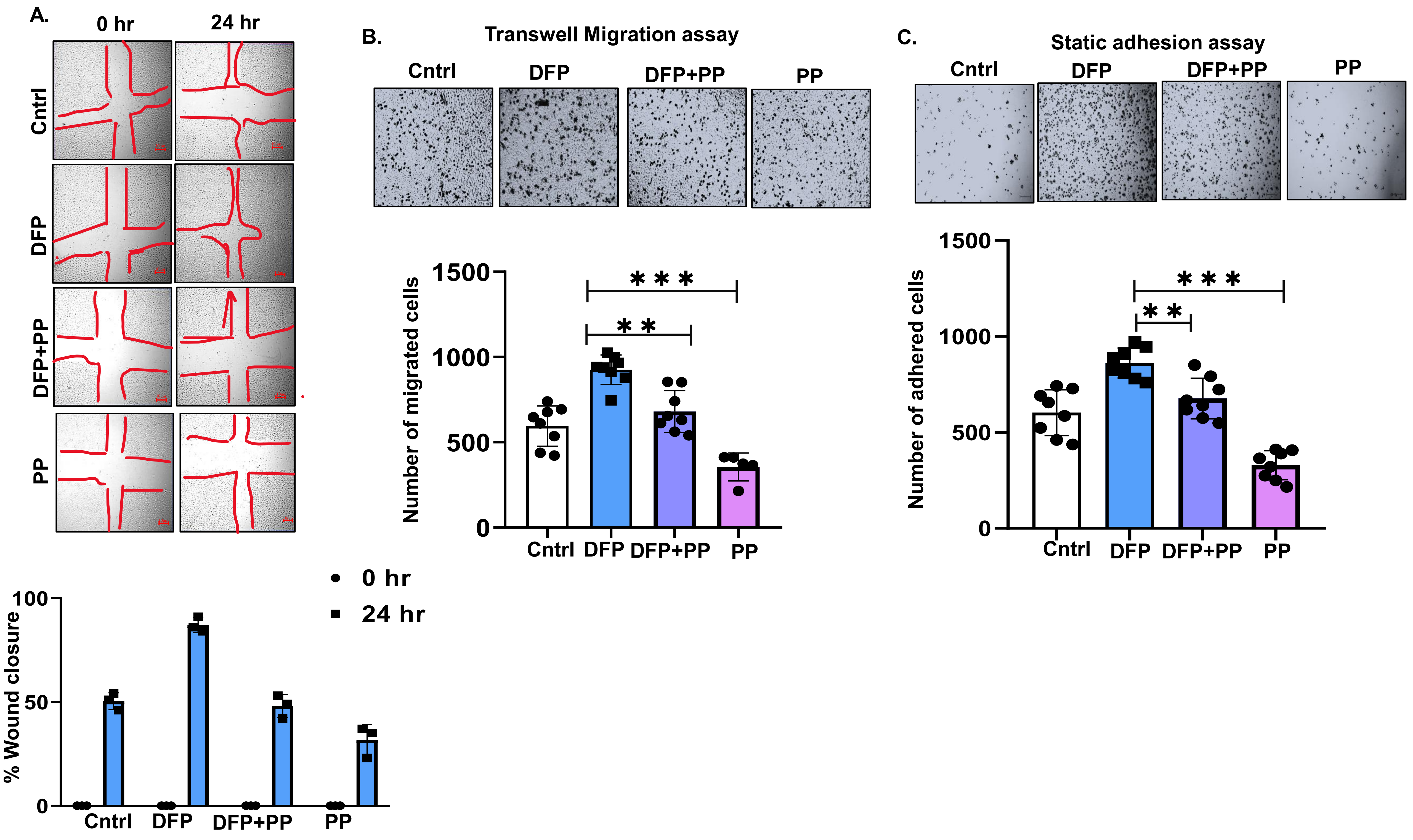

**Supplementary Figure 7**

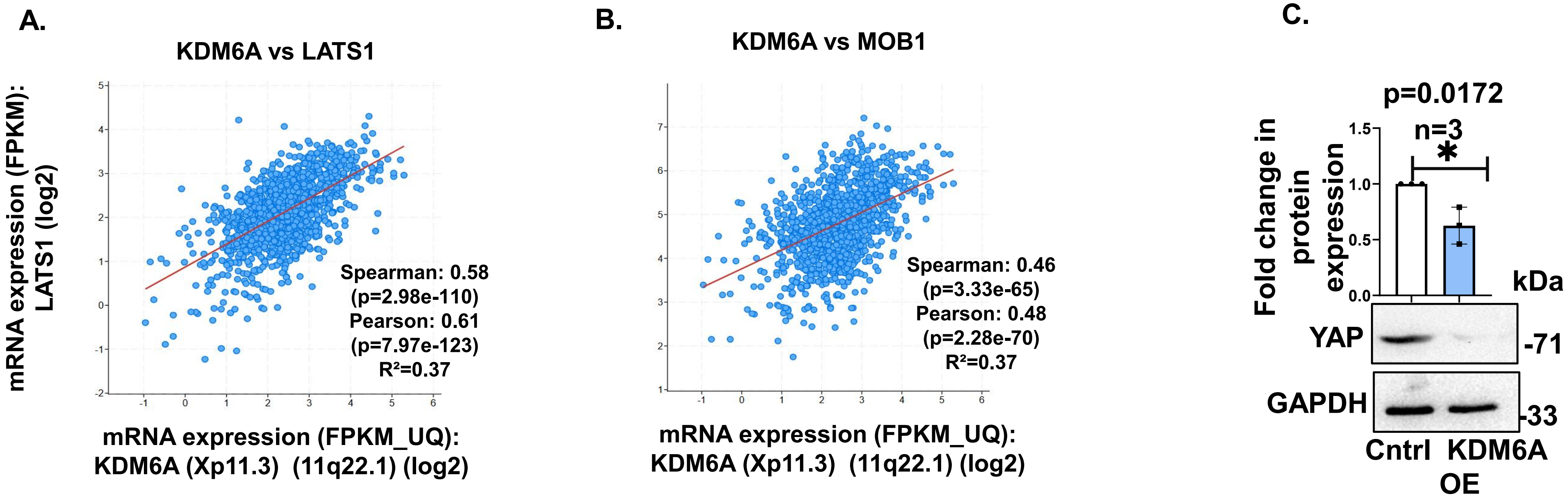

Supplementary Figure 8

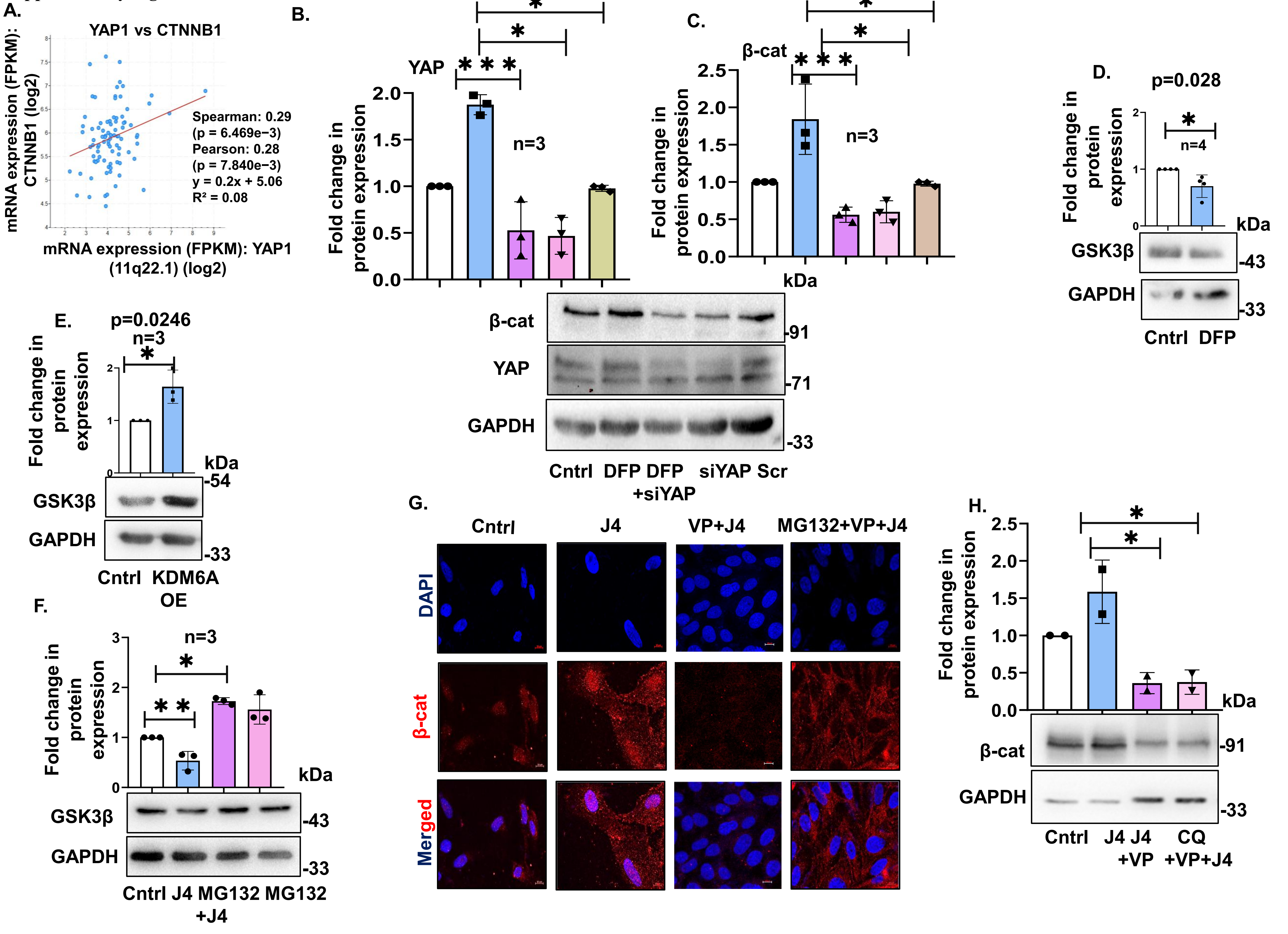
